# Structural basis for substrate specificity and MSMEG_0435-0436 binding by the mycobacterial long-chain acyl-CoA carboxylase complex

**DOI:** 10.1101/2025.10.28.685139

**Authors:** Yingke Liang, John L. Rubinstein

## Abstract

The presence of mycolic acid is a defining feature of the mycobacterial cell wall, which provides a highly impermeable barrier to many antibiotics. Biosynthesis of this fatty acid, as well as tuberculostearic acid, requires precursor molecules produced by the essential long-chain acyl-coenzyme A (CoA) carboxylase (LCC) complex. The LCC complex catalyzes carboxylation of the α-carbon of long-chain acyl-CoA, but also short-chain acetyl-CoA and propionyl-CoA. The complex includes the subunits AccA3, which contains a biotin carboxylase (BC) domain and a biotin carboxyl carrier protein (BCCP) domain, the long-chain acyl-CoA carboxyltransferase AccD4, the short-chain acyl-CoA carboxyltransferase AccD5, and the incompletely characterized protein AccE. We used electron cryomicroscopy (cryo-EM) to determine structures of the LCC complex from *Mycobacterium smegmatis*. In the structures, two AccE subunits tether eight AccA3 subunits to an AccD4_2_AccD5_4_ heterohexamer core. Cryo-EM of the enzyme during catalysis reveals how AccD4 and AccD5 achieve substrate specificity, with AccD5 binding tightly to CoA and AccD4 binding long acyl chains. The BCCP domains of AccA3 undergo long-distance translocation to transfer a carboxyl group from the BC domain of AccA3 to the acyl-CoA substrate bound in AccD5. Further, we find that two copies of a protein complex formed from MSMEG_0435 and MSMEG_0436 can bind the LCC complex, sequestering the biotin moiety of BCCP domains near AccD5. Rv0263c, the *Mycobacterium tuberculosis* ortholog of MSMEG_0435, has a role in bacterial survival during transmission, suggesting that these proteins may regulate production of branched fatty acid precursors for the mycobacterial cell wall.

## Introduction

A defining characteristic of mycobacteria and other members of the suborder *Corynebacterineae* is a cell wall rich in mycolic acids (Marrakchi et al., 2014; Stodola et al., 1938; Zuber et al., 2008). These fatty acids have a long (54 to 63 carbon) acyl chain with a hydroxyl group on their β-carbon, and a shorter (22 to 24 carbon) saturated alkyl side chain branching from their α-carbon. The long-chain fatty acid, known as meromycolate, has characteristic modifications including cyclopropane rings and methoxy or keto groups. Mycolic acids form the mycomembrane layer of the cell wall, with this layer comprising an inner leaflet of mycolic acid linked to arabinogalactan and an outer leaflet thought to consist of free trehalose monomycolate, trehalose dimycolate, and other glycolipids (Marrakchi et al., 2014; Zuber et al., 2008). For pathogenic mycobacteria, including *Mycobacterium tuberculosis*, the etiologic agent of the disease tuberculosis (TB), the mycomembrane offers protection against antibiotics and the host immune system.

Synthesis of mycolic acid, along with 10-methyloctadecanoic acid or tuberculostearic acid found in the mycobacterial plasma membrane (Daffé and Marrakchi, 2019), requires acyl-coenzyme A (acyl-CoA) precursors that are carboxylated at their α-carbon (Marrakchi et al., 2014). These precursors are produced by biotin-dependent acyl-CoA carboxylases (Tong, 2013). Mycobacterial acyl-CoA carboxylases are formed from α and β subunits: the α subunit includes a biotin carboxylase (BC) domain and a biotin carboxyl carrier protein (BCCP) domain, while the β subunit contains a carboxyltransferase (CT) domain. The BC domain uses bicarbonate and energy from adenosine triphosphate (ATP) hydrolysis to transfer a carboxyl group to a biotinylated lysine residue from the BCCP domain. The CT domain binds the acyl-CoA substrate, thereby determining substrate specificity, and transfers the carboxyl group from the carboxybiotin of the BCCP domain to the α-carbon of the acyl-CoA.

Mycobacterial genomes encode an unusually large number of biotin-dependent acyl-CoA carboxylases, not all of which appear to be involved in lipid metabolism. Three genes (*accA1* to *accA3*) encode α subunits and six (*accD1* to *accD6*) encode β subunits (Ehebauer et al., 2015).

An ε subunit, encoded by *accE* (sometimes called *accE5*), has an unclear function but is found in some acyl-CoA carboxylases in actinobacteria, including mycobacteria (Gago et al., 2006). The genes *accA3*, *accD4*, and *accD5* appear essential in *M. tuberculosis* by both transposon insertion sequencing (TnSeq) (DeJesus et al., 2017) and CRISPRi screening (Bosch et al., 2021). The gene *accD6* has a growth defect by TnSeq but appears to be essential by CRISPRi screening, while *accE* appears essential by TnSeq but not CRISPRi screening. The genes *accA1*, *accA2*, *accD1*, *accD2*, and *accD3* are not essential.

The AccA1 and AccA2 proteins form complexes with AccD1 and AccD2, respectively (Ehebauer et al., 2015). The AccA1-AccD1 complex carboxylates 3-methylcrotonyl-CoA during leucine degradation and is known as the 3-methylcrotonyl-CoA carboxylase (MCC) complex. Its close paralog, the AccA2-AccD2 complex, probably has the same activity (Ehebauer et al., 2015). AccD3 does not appear to interact with an AccA protein (Ehebauer et al., 2015) and may not include a CT domain at all (Gande et al., 2004). AccA3 interacts with AccD4, AccD5, and likely AccD6 (Ehebauer et al., 2015). AccD4 carboxylates long-chain acyl-CoA (Oh et al., 2006; Portevin et al., 2005), while AccD6 catalyzes carboxylation of acetyl-CoA (Daniel et al., 2007; Kurth et al., 2009; Pawelczyk et al., 2017, 2011), and AccD5 carboxylates both propionyl-CoA and acetyl-CoA (Gago et al., 2006). An essential complex formed from AccA3, AccD4, AccD5, and AccE known as the long-chain acyl-CoA carboxylase (LCC) complex, can carboxylate both acetyl- and propionyl-CoA with the AccD5 subunit and long-chain acyl-CoA with the AccD4 subunit (Bazet Lyonnet et al., 2017). A long-chain acyl-CoA carboxylated at its α-carbon is needed to form the α branch of mycolic acids (Oh et al., 2006), while malonyl-CoA, the product of acetyl-CoA carboxylation by AccD4 (and AccD6), is a substrate for the type II fatty acid synthesis system in mycobacteria used for producing the main meromycolate chain of mycolic acid (Marrakchi et al., 2014). Methylmalonyl-CoA, the product of propionyl-CoA carboxylation by AccD5, likely serves as a precursor for tuberculostearic acid in the mycobacterial plasma membrane (Daffé and Marrakchi, 2019).

We inadvertently found that Strep-Tactin XT affinity matrix can be used to isolate endogenously biotinylated proteins from *Mycobacterium smegmatis*. Electron cryomicroscopy (cryo-EM) of these proteins allowed us to determine the structure of the LCC complex and the MCC complex. The MCC complex has the stoichiometry AccA1_6_AccD1_6_ and closely resembles structures of the human and *Pseudomonas aeruginosa* MCC complexes (Huang et al., 2011; Su et al., 2025). The LCC complex has a stoichiometry of AccA3_8_AccD4_2_AccD5_4_AccE_2_. In the structure, a central AccD4_2_AccD5_4_ heterohexamer is flanked on each face by two pairs of AccA3 subunits, with each of two AccE subunits responsible for tethering four AccA3 subunits. The BCCP and BC domains of an AccA3 subunit contact the AccD4 subunits, but the BCCP domain would need to rotate by ∼180° for long-chain acyl-CoA carboxylation. In contrast, each of the AccD5 subunits has a BCCP domain from an AccA3 subunit bound and oriented correctly for short-chain acyl-CoA carboxylation. However, the BC domain of the AccA3 subunit is too far away to transfer a carboxyl group to the biotin of the BCCP domain. Structure determination of the LCC complex in the presence of substrate shows how AccD5 selectively binds a short-chain acyl-CoA while AccD4 binds a long-chain acyl-CoA. Under turnover conditions, the BCCP domains bound to the AccD5 subunits become mobile, enabling observation of a different conformation where the BCCP domain interacts with the BC domain to receive the carboxyl group. Finally, we identified a subpopulation of particle images where complexes of MSMEG_0435 and MSMEG_0436 are bound to the AccD5 subunits and BCCP domains from AccA3 subunits. Each bound MSMEG_0435 displaces one BCCP domain and sequesters the biotin moiety from another. MSMEG_0435 and MSMEG_0436 are orthologs of *M. tuberculosis* Rv0263c and Rv0264c, with Rv0263c recently found to be necessary for *M. tuberculosis* survival in a model of TB transmission (Singh et al., 2025).

## Results

### Isolation of the LCC and MCC complexes from M. smegmatis

When using Strep-Tactin XT affinity matrix and size exclusion chromatography to isolate Twin-Strep-tagged proteins from *M. smegmatis*, we found that two large protein complexes contaminated cryo-EM images (**Fig. S1A**, *purple* and *yellow* boxes). *Ab initio* three-dimensional (3D) reconstruction (Punjani et al., 2017) followed by map refinement and automated backbone modeling (Jamali et al., 2024) allowed construction of initial atomic models for these complexes. These models were used to search a database of protein folds (Van Kempen et al., 2024), enabling identification of the complexes as the MCC complex, formed from subunits AccA1 and AccD1, and the LCC complex, including subunits AccA3, AccD4, AccD5, and AccE. Mass spectrometry confirmed the presence of these proteins in the sample used for cryo-EM (**Supplementary Data 1**). AccA2, AccD2, and pyruvate carboxylase were also identified in the sample by mass spectrometry but did not give rise to cryo-EM maps. In mycobacteria, the AccA subunits (**Fig. 1A**, *blue rectangle*) contain the BC and BCCP domains, while the AccD subunits (**Fig. 1A**, *pink and orange rectangle*) contain the CT domain. Refinement with D3 symmetry imposed allowed for calculation of a 2.0 Å resolution map of the MCC complex (**Fig. 1B**, *left*, **Fig. S1**, **Table S1**), and allowed for construction of an atomic model for 93% of residues in the complex (**Fig. 1B**, *top right*, **Table S4**). The map of the LCC complex could be refined to an overall nominal resolution of 2.9 Å (**Fig. S2**, **Table S2**), but the flexible AccA3 subunits required local refinement to reach high resolution (**Fig. S3**). A composite map of the LCC complex (**Fig. 1C**, *left*) allowed construction of an atomic model for 86% of residues in the LCC complex (**Fig. 1C**, *right,* **Fig. 1D**, **Table S4**).

**Figure 1.**
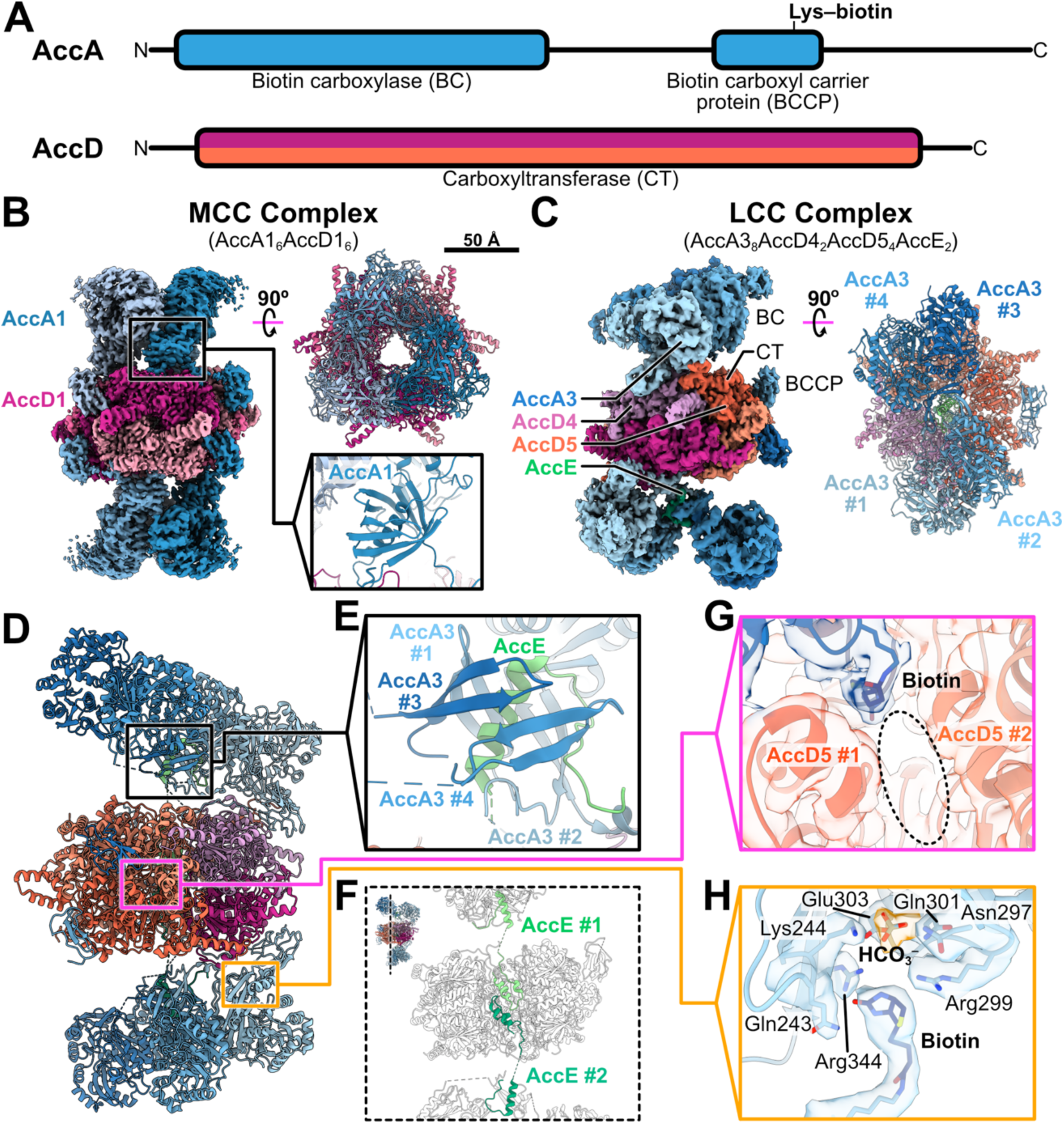
Overall architecture of the mycobacterial MCC complex and LCC complex. **A,** Domain architecture for the AccA (*upper*) and AccD (*lower*) subunits of the 3-methylcrotonyl-CoA carboxylase (MCC) and long-chain acyl-CoA carboxylase (LCC) complexes. **B,** Cryo-EM map (*left*) and atomic model (*right*) for the MCC complex. **C,** Composite cryo-EM map (*left*) and a top view of the atomic model (*right*) for the LCC complex. BC, biotin carboxylase domain. BCCP, biotin carboxyl carrier protein domain. CT, carboxyltransferase domain. **D**, A side view of the atomic model for the LCC complex. **E,** The N-terminal α helix from the AccE subunit of the LCC complex binds β hairpins from four AccA3 subunits. **F,** A C-terminal α-helical hairpin anchors AccE to the rest of the LCC complex. **G,** When AccA3 is bound to an AccD5 subunit, the biotin moiety from its BCCP domain is resolved at the interface between two AccD5 subunits. The dashed oval indicates the short-chain acyl-CoA binding site. **H,** When AccA3 is bound to an AccD4 subunit, the biotin moiety from its BCCP domain interacts with its BC domain.

### The MCC complex architecture resembles other known MCC structures

The MCC complex plays a role in mycobacterial leucine catabolism (Ehebauer et al., 2015) and its structure determined here resembles structures of human and *Pseudomonas aeruginosa* MCC complexes (Huang et al., 2011; Su et al., 2025). In the structure, which has D3 symmetry, six AccD1 subunits form a doughnut-shaped hexamer comprising two stacked rings of AccD1 trimers (**Fig. 1B**, *pink)*. Three AccA1 subunits extend away from each of the two flat surfaces of the hexamer (**Fig. 1B**, *blue*). The N-terminal BC domain of each AccA1 subunit is attached by an extended linker to a domain comprising an eight-stranded β barrel with a single α helix in the middle of the barrel, which anchors the BC domain onto an AccD1 subunit (**Fig. 1B**, *bottom right*). An additional extended linker connects this anchoring domain to the BCCP domain of AccA1, which is tethered at the periphery of an adjacent AccD1 subunit. Similar β barrels perform the same function in other known MCC complex structures (Huang et al., 2011; Su et al., 2025) and propionyl-CoA carboxylase structures (Huang et al., 2010; Lee et al., 2023; Scheffen et al., 2021; Zhou et al., 2024). Because the structure of the MCC complex has been described extensively from other organisms, we focused our analysis instead on the LCC complex, the structure of which has not been described previously.

### The mycobacterial LCC complex architecture differs from other acyl-CoA carboxylases

The structure of the LCC complex does not resemble other known acyl-CoA carboxylase complexes. Instead, the LCC complex consists of a heterohexameric core of two AccD4 subunits (**Fig. 1C** and **D**, *pink*) and four AccD5 subunits (**Fig. 1C** and **D**, *orange*). Like the AccD1_6_ hexamer of the MCC complex, this AccD4_2_AccD5_4_ hexamer resembles a doughnut made from two stacked AccD4AccD5_2_ heterotrimer rings. Because each half of the heterohexamer is a heterotrimer rather than a homotrimer, the AccD4_2_AccD5_4_ hexamer has only C2 symmetry with pseudo-D3 symmetry. AccE subunits (**Fig. 1C** to **F**, *green*) are tethered by C-terminal α-helical hairpins to the center of the AccD4_2_AccD5_4_ core (**Fig. 1F**), with each extending in opposite directions from the flat surfaces of the hexamer. The N-terminal α helices from each AccE subunit (**Fig. 1E**, *green*) bind four AccA3 subunits (**Fig. 1C** to **E**, *blue*), anchoring them to each side of the AccD4_2_AccD5_4_ hexamer and resulting in eight AccA3 protomers per LCC complex. Overall, the LCC complex has a subunit composition of AccA3_8_AccD4_2_AccD5_4_AccE_2_.

However, other 3D classes from the dataset show AccD4_2_AccD5_4_ hexamers with AccA3 subunits only on one side of the hexamer or with only the AccD4_2_AccD5_4_ hexamer resolved, almost certainly because of flexibility in the positions of the AccA3 subunits (**Fig. S4A**).

The AccA3 subunits each include a BC domain and BCCP domain (**Fig. 1A**, *top*, **Fig. 1C**, *blue*). The linker sequences that connect the BC and BCCP domains are generally not resolved in the structure, but β hairpins in the middle of the linkers are resolved where they interact with AccE (**Fig. 1E**, *blue*). These β hairpins from four AccA3 subunits form an eight stranded β barrel around an α helix from the AccE subunit (**Fig. 1E**, *green*) that resembles the structures that link the BC and BCCP domains of the AccA1 and tether the BC domain of AccA1 to AccD1 in the MCC complex (**Fig. 1B**, *bottom right*). While BC domains and β hairpins from the linker sequence are resolved for all eight AccA3 subunits in the complex, only six of the BCCP domains are resolved. These six BCCP domains are bound to the six protomers in the AccD4_2_AccD5_4_ hexamer. The eight AccA3 subunits for two AccD4 and four AccD5 subunits is remarkable because it means there are more BC and BCCP domains available in the complex to produce carboxybiotin than there are CT domains for carboxylation of substrate. In contrast, other known carboxyltransferase structures have an equal number of BC, BCCP, and CT domains.

### AccD4 and AccD5 interact differently with the BCCP and BC domains of AccA3

The BCCP domains bound to the AccD5 subunits (**Fig. 1C** and **D**, *blue* and *orang*e) interact in a manner comparable to the interaction between the BCCP and CT domains in other carboxylase enzymes (Diacovich et al., 2004; Huang et al., 2011, 2010; Scheffen et al., 2021; Su et al., 2025; Zhou et al., 2024). In this interaction, the biotin moiety from the BCCP extends into an unobstructed groove in AccD5 close to the putative short-chain acyl-CoA binding site **(Fig. 1G**, *dashed oval*). The BCCP domains bound to the AccD5 subunits do not contact a BC domain. Owing to the flexible linker between the BCCP and BC domains of these AccA3 subunits, it is not possible to say with complete confidence which BC domain and which BCCP domain are from the same polypeptide. In contrast, the AccA3 and AccD4 subunits (**Fig. 1C** and **D**, *blue* and *pink*) interact in a non-canonical manner. Each AccD4 subunit binds both the BCCP and BC domain of an AccA3 subunit. However, the BCCP domain bound to AccD4 is rotated by ∼180° compared to how these domains interact with AccD5, so that the biotin points into the BC domain of the AccA3 (**Fig. 1H**). Within this BC domain, a flattened density in the cryo-EM map ∼5 Å from the biotin, bound near basic and polar residues, could be modeled as a bicarbonate (**Fig. 1H**, *orange density*). As a result of the close interaction between its BCCP and BC domain, the entirety of this AccA3 subunit could be modelled. AccD4 also interacts with residues 511 to 517 of AccA3 through multiple polar interactions (**Fig. S4B**). This interaction is visible even when the AccA3 cannot be resolved and may be important for maintaining an AccA3 close to AccD4 for catalysis.

### Structural basis for short-chain acyl-CoA carboxylase activity

The LCC complex is expected to possess long-chain acyl-CoA, acetyl-CoA, and propionyl-CoA carboxylase activity, owing to the presence of both AccD4 and AccD5 in the assembly (Gago et al., 2006; Oh et al., 2006; Portevin et al., 2005). We used an enzyme-coupled assay (Janiyani et al., 2001) with ATP, sodium bicarbonate, and propionyl-CoA as substrates to confirm that the protein preparation has short-chain acyl-CoA carboxylase activity (**Fig. 2A**, **Fig. S5**). To investigate the structural basis for this activity, we prepared cryo-EM grids of the sample in the presence of the three substrates (**Fig. S6**). *Ab initio* 3D reconstruction followed by iterative refinement again allowed calculation of a map of the LCC complex with a nominal overall resolution of 2.3 Å, with local refinement improving map density for the AccA3 subunits (**Fig. 2B**, **Fig. S7**, **Table S3**). A composite map enabled construction of an atomic model for 94% of residues in the complex (**Table S4**). Within the BC domains, residues 140-299 of each AccA3 subunit form an ATP grasp fold responsible for nucleotide binding (Fawaz et al., 2011).

**Figure 2.**
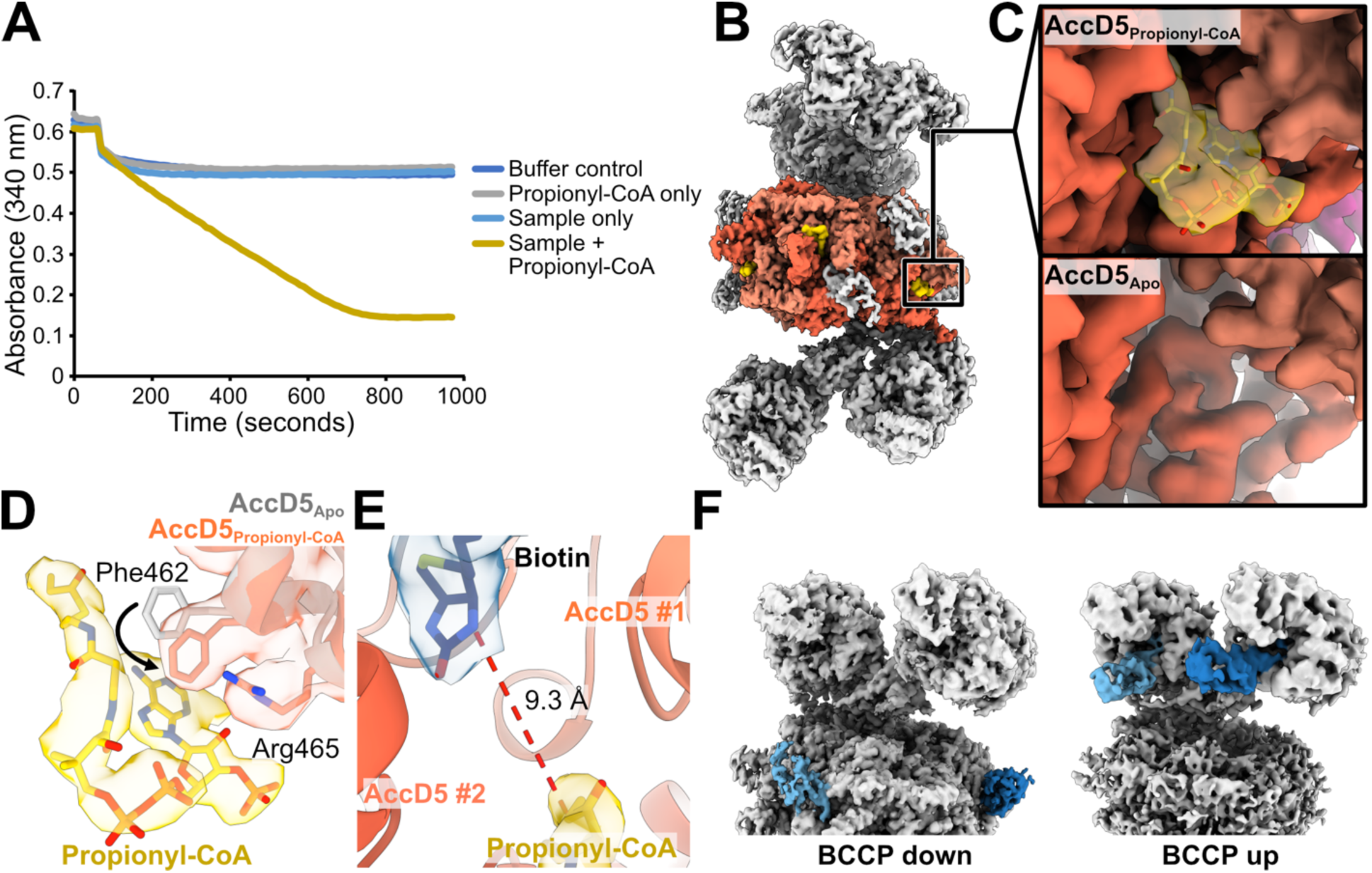
Substrate binding and BCCP domain movement for carboxylation of short-chain fatty acids by the LCC complex. **A,** The sample displays robust ATPase activity when provided with propionyl-CoA, bicarbonate, and ATP. **B,** Composite cryo-EM map of the LCC complex with propionyl-CoA bound. **C,** Density for propionyl-CoA (*yellow*) in the AccD5 subunit is seen in the presence of substrate (*upper*) but not the absence of substrate (*lower*). **D,** Binding of propionyl-CoA to AccD5 induces movement of Phe462 and stabilization of Arg465. **E,** Bound propionyl-CoA in AccD5 is too far from the biotin moiety (9.3 Å) for transfer of a carboxyl group. **F,** Incubating the LCC complex with substrate causes AccA3 BCCP domains (*blue*) to translocate from “down” conformations interacting with an AccD5 subunit to “up” conformations interacting with the AccA3 BC domain.

Consistent with previous structures of ATP grasp folds, density was absent for residues 140-208, which form the lid, in the map of the LCC complex obtained without ATP (**Fig. S8A**, *top right*) but was better ordered in the presence of ATP (**Fig. S8A**, *bottom right*).

Propionyl-CoA is bound only to the AccD5 subunits and not the AccD4 subunits (**Fig. 2B** and **C**, *yellow density*) and is well resolved in all acyl-CoA binding pockets in AccD5. Binding of propionyl-CoA to AccD5 induces movement of Phe462, positioning it to form a π–π interaction with the purine moiety of CoA, while Arg465 adopts a more extended conformation where it interacts with the 2′ hydroxyl of the ribose moiety of CoA (**Fig. 2D**). In the cryo-EM map, the interaction of the BCCP domain with AccD5 appeared unchanged from the map obtained without substrate. In this conformation, the biotin moiety from the BCCP domain is ∼8 Å from the propionyl-CoA (**Fig. 2E**) and there is no evidence for carboxylation of the biotin.

However, diffuse density extending from the BC domains of two of the AccA3 subunits near the AccD5 subunits suggested conformational heterogeneity in the dataset (**Fig. S8B**, *red density*). Three-dimensional classification without image alignment, focusing on the region of the map adjacent to these BC domains, allowed separation of the particle images into four classes. In one class, the BCCP domains associated with the AccD5 subunits remain bound to the AccD5 subunits in “down” conformations (**Fig. 2F**, *left - blue densities*). In another, both BCCP domains associated with the AccD5 subunits no longer contact AccD5 and instead contact the BC domain of AccA3 in “up” conformations (**Fig. 2F**, *right - blue densities*). In the remaining two classes, only one BCCP domain is in the “up” conformation. The existence of these classes indicates that the BCCP domains of AccA3 independently traverse the large distance from the AccD5 subunit to the BC domain of the AccA3 subunit (**Movie 1**). This movement of the BCCP domains could only be detected in the dataset with substrate added, with the “up” conformation of the BCCP domain likely showing a state ready for biotin carboxylation by the BC domain.

However, it is possible that the BCCP may translocate to the BC in the absence of added substrate at a much lower rate that is difficult to capture by cryo-EM. Substrate-induced translocation of a BCCP domain has been observed previously in pyruvate carboxylases (Chai et al., 2022). In contrast, in MCC complexes the BCCP domain alternates between the BC- interacting and CT-interacting conformations even in the absence of substrate (Su et al., 2025).

### Structural basis for long-chain acyl-CoA carboxylase activity

The *in vivo* substrates for AccD4 in mycobacteria are thought to be CoA esters of fatty acids with between 24 and 26 carbons (Oh et al., 2006; Portevin et al., 2005). However, *in vitro*, AccD4 can carboxylate acyl-CoA with as few as 16 carbons in the acyl chain (Bazet Lyonnet et al., 2017). The differential specificity of AccD4 and AccD5 for long- and short-chain acyl-CoA substrates, respectively, occurs despite them having a conserved overall fold with 63% amino acid sequence similarity and 49% sequence identity. To confirm that the enzyme preparation has long-chain acyl-CoA carboxylase activity, we repeated the enzyme coupled assay but replaced propionyl-CoA with arachidoyl-CoA, which has a saturated 20-carbon acyl chain (**Fig. 3A**, **Fig. S5B**). This experiment revealed robust activity, although ∼60× slower than the propionyl-CoA carboxylase activity at an equivalent concentration of protein. To understand how the LCC complex binds both short-chain and long-chain acyl-CoA substrates, we prepared cryo-EM specimens of the protein preparation with both arachidoyl-CoA and propionyl-CoA, as well as ATP and bicarbonate (**Fig. S9**). *Ab initio* 3D reconstruction followed by iterative refinement allowed calculation of a 2.0 Å resolution map of the complex, again requiring local refinement to improve map density for the AccA3 subunits (**Fig. 3B**, **Table S1**, **Fig. S10**). A composite map prepared from this map and locally refined maps of the AccA3 subunits enabled construction of an atomic model for 94% of residues in the complex (**Table S4**).

**Figure 3.**
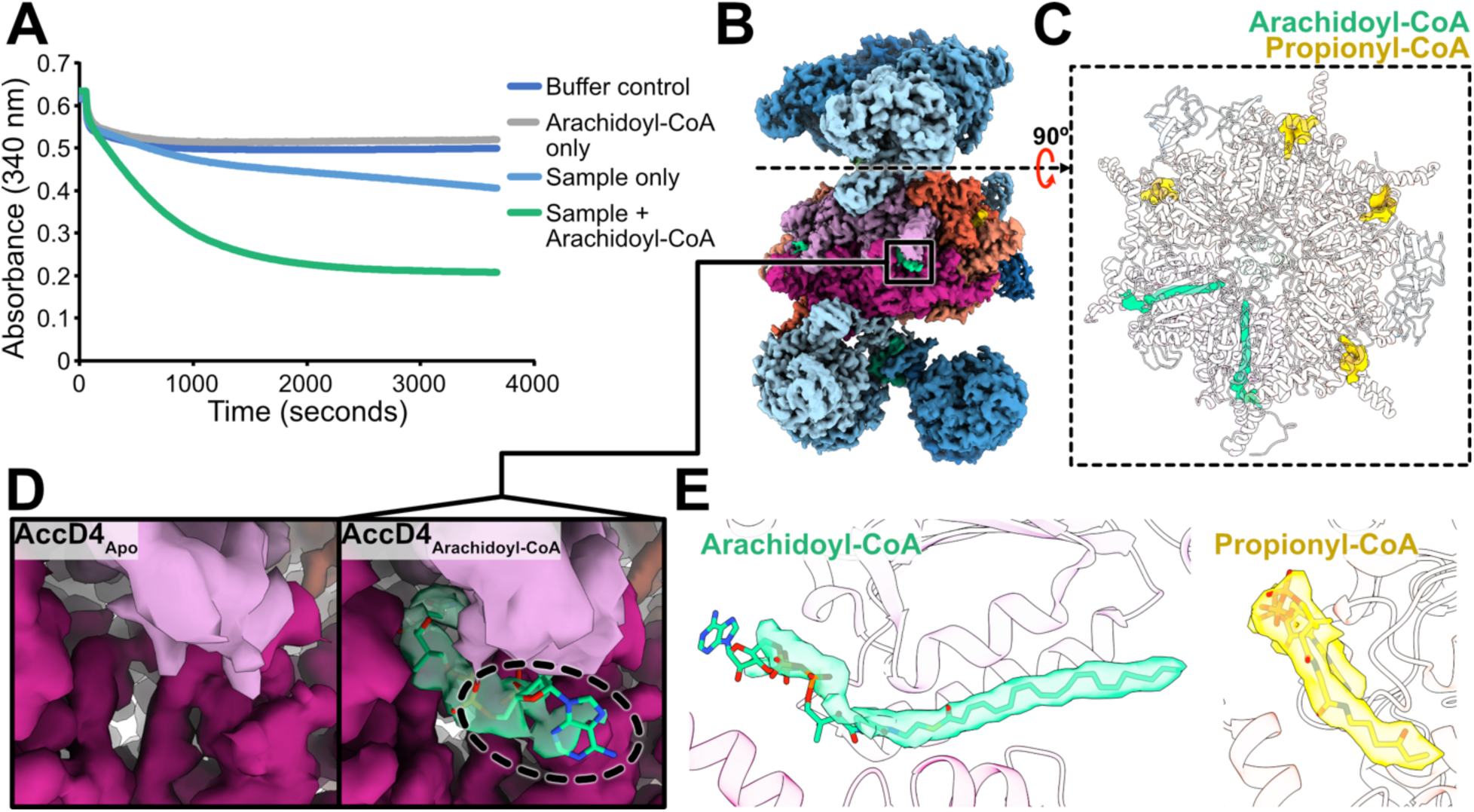
Substrate binding for carboxylation of long-chain fatty acids by the LCC complex. **A,** The sample displays ATPase activity in the presence of arachidoyl-CoA with bicarbonate and ATP, but with ∼60× lower activity than with propionyl-CoA at an equivalent concentration of protein. **B,** Composite cryo-EM map of the LCC complex in the presence of both short- and long-chain acyl-CoA substrates shows density for propionyl-CoA (*yellow*) and arachidoyl-CoA (*green*). **C,** Cross-section through the atomic model showing the propionyl-CoA and arachidoyl-CoA binding sites. **D,** Comparison of the long-chain acyl-CoA binding site in AccD4 without (*left*) and with (*right*) arachidoyl-CoA added. The CoA moiety of arachidoyl-CoA is poorly resolved (*dashed oval*). **E,** Comparison of the cryo-EM map densities for arachidoyl-CoA (*left*) and propionyl-CoA (*right*) shows good resolution for the acyl tail of arachidoyl-CoA and the entire propionyl-CoA molecule.

Overall, the map resembles maps of the LCC complex without substrate or bound to propionyl-CoA, but density is present for both arachidoyl-CoA and propionyl-CoA. Arachidoyl-CoA is bound in the AccD4 subunits (**Fig. 3C**, *magenta densities*), in a pocket that resembles where propionyl-CoA binds to AccD5 (**Fig. 3C**, *yellow densities*). The structure of AccD4 creates an elongated tunnel that allows the acyl chain of arachidoyl-CoA to penetrate the AccD4_2_AccD5_4_ hexamer (**Fig. 3D**). The CoA moiety from arachidoyl-CoA is poorly resolved (**Fig. 3D**, *right – dashed oval*), indicating that this part of the molecule remains loosely bound. In contrast, the acyl chain of arachidoyl-CoA is well resolved, indicating that AccD4 binds it tightly (**Fig. 3E**, *left*). This tight binding of long acyl chains by AccD4 is consistent with kinetic measurements that showed that the LCC complex has a higher affinity for long-chain acyl-CoA than propionyl- or acetyl-CoA, as indicated by a lower Michaelis constant (Bazet Lyonnet et al., 2017). Whereas the entirety of propionyl-CoA is resolved well in the map (**Fig. 3E**, *right*), AccD4 lacks an equivalent of the Phe462 residue found in AccD5, which stabilizes the interaction of the CoA moiety. These observations suggest that short-chain acyl-CoA binding to AccD5 occurs through specific interactions with the CoA moiety, while long-chain acyl-CoA binding in AccD4 is achieved through interactions with the long acyl chain. Reliance on different parts of the acyl-CoA for binding and AccD5 lacking the ability to accommodate a long-chain acyl-CoA likely explains why the two subunits possess different specificities for long- and short-chain acyl-CoA. In the structure, the position of the BCCP domain near AccD4 remains unchanged. This conformation places the biotin moiety from the BCCP domain close to the BC domain, in a state where the biotin is poised for carboxylation by the BC but relatively far from the arachidoyl-CoA substrate.

### An MSMEG_0435-MSMEG_0436 complex binds the LCC complex

In the dataset of particle images frozen with arachidoyl-CoA, propionyl-CoA, ATP, and bicarbonate (**Table S1**), we identified 3D classes of particle images that produced maps with one or two extra densities. These regions of the map were refined to 2.9 Å resolution, and automated backbone modeling (Jamali et al., 2024) and search of a database of protein folds (Van Kempen et al., 2024) were again used to identify the proteins corresponding to the densities. This pipeline identified the additional proteins as MSMEG_0435 and MSMEG_0436, which were also identified in mass spectrometry of the sample (**Supplementary Data 1**). The structures show either one or two copies of the MSMEG_0435-MSMEG_0436 complex, each bound to an AccD5 subunit and two BCCP domains from AccA3 subunits (**Fig. 4A**, *purple*, **Fig. S10**). An atomic model of MSMEG_0435-MSMEG_0436 interacting with the LCC was built from the cryo-EM map with two copies of the complex bound and allowed for modelling of 96% of residues in the complex (**Fig. 4A**, *right*, **Table S4**). MSMEG_0435 binds the AccD5 subunits while MSMEG_0436 is distal from the AccD4_2_AccD5_4_ hexamer and does not interact with it directly. Remarkably, whereas only six of the BCCP domains from the eight AccA3 subunits were resolved in the structures described above, all eight BCCP domains are resolved in the 3D classes where two of the MSMEG_0435-MSMEG_0436 complexes are bound. The additional BCCP domains are resolved near each pair of AccD5 subunits on the two sides of the AccD4_2_AccD5_4_ heterohexamer core (**Fig. 4B**). The BCCP domains associated with the AccD4 subunit remain unperturbed. Each MSMEG_0435-MSMEG_0436 complex occludes a surface on AccD5 that was previously occupied by a BCCP domain (**Fig. 4C**, *left*). Binding of the MSMEG_0435-MSMEG_0436 complex displaces this BCCP domain and the displaced BCCP domain, along with an additional BCCP domain, bind a new site on MSMEG_0435 (**Fig. 4C**, *right*). In one of the two BCCP domains bound to MSMEG_0435 the biotin moiety and associated lysine can no longer be resolved in the cryo-EM map, suggesting that they are disordered in the structure. Further, each MSMEG_0435 protein appears to bind tightly to the biotin moiety of the other BCCP domain (**Fig. 4D**). These observations suggest that neither BCCP domain that interacts with MSMEG_0435 would function with AccD5. A single BCCP domain remains bound to AccD5 on each side of the AccD4_2_AccD5_4_ hexamer, maintaining the interaction needed for carboxylation of short-chain acyl-CoA. MSMEG_0435 is homologous with urea carboxylases, and the biotin from the additional BCCP is positioned to participate in a carboxylation reaction within the protein. However, several residues important for urea carboxylase activity are absent from the protein sequence (Fan et al., 2012). Using an enzyme coupled assay that has been used previously to measure urea carboxylase activity (Kanamori et al., 2004; Lin et al., 2016) we confirmed that the protein preparation does not possess this activity, even at 350 µg/mL total protein (**Fig. S5C**).

**Figure 4.**
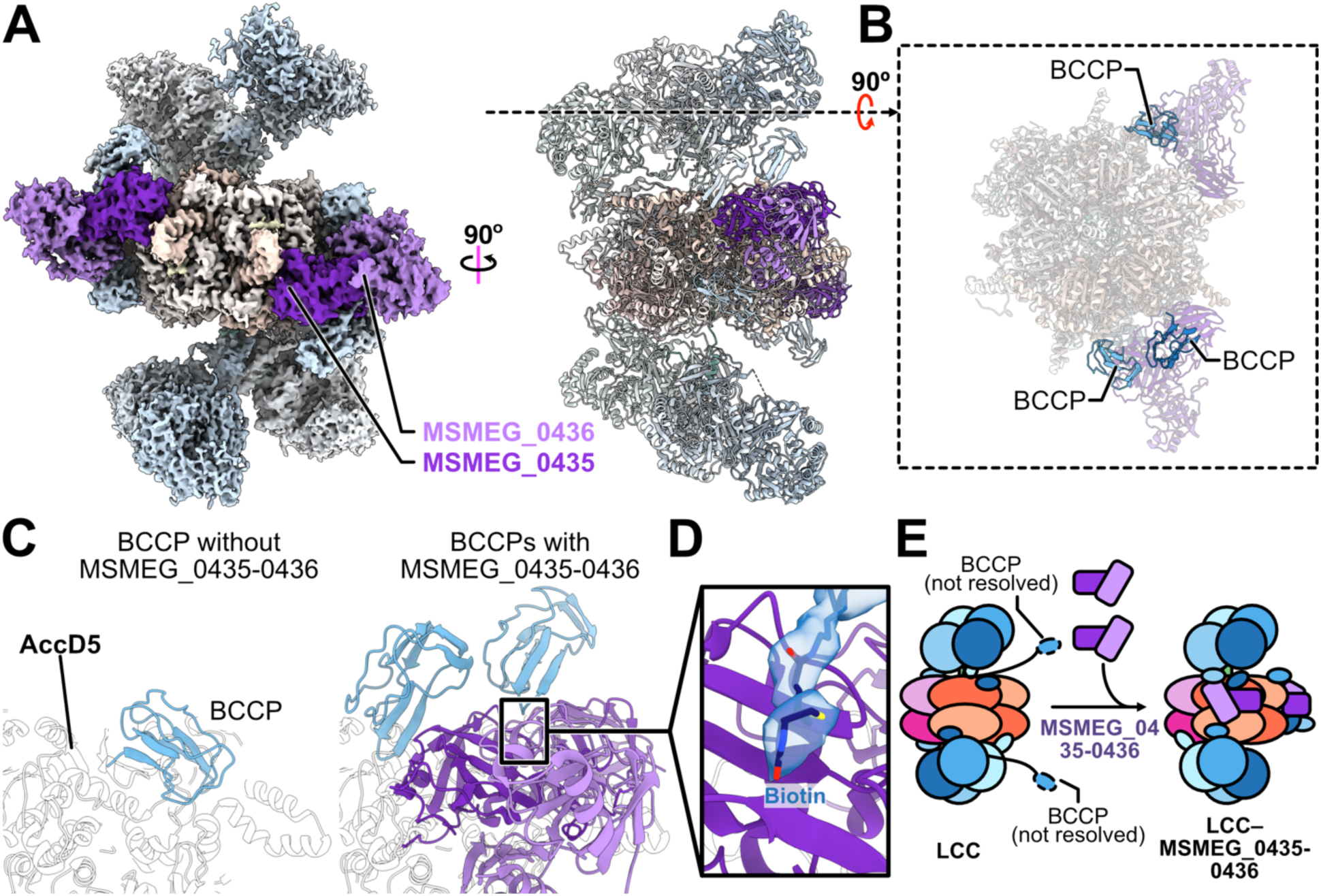
MSMEG_0435 and MSMEG_0436 bind the LCC complex and sequester biotin from AccA3. **A,** Cryo-EM map (*left*) and atomic model (*right*) for the LCC complex bound to two copies of the MSMEG_0435-MSMEG_0436 complex (*purple*). **B,** Two additional BCCP domains from the AccA3 subunits (*blue*) are resolved near AccD5 subunits in the structure when two MSMEG_0435-MSMEG_0436 complexes are bound. **C,** MSMEG_0435-MSMEG_0436 binding displaces a BCCP domain from its normal position on AccD5 (*left*) and binds the BCCP domain and an additional BCCP domain (*right*). **D,** MSMEG_0435 sequesters a biotin moiety from one of the BCCP domains. **E,** Schematic of conformational changes in the LCC complex on MSMEG_0435-MSMEG_0436 binding.

## Discussion

We determined structures of both the MCC complex and the LCC complex from *M. smegmatis*. The MCC complex structure appears similar to MCC complexes from other organisms. To our knowledge, there are no available structures of other LCC complexes. The LCC complex catalyzes the carboxylation of acetyl- and propionyl-CoA with its AccD5 subunits, and the carboxylation of long-chain acyl-CoA with its AccD4 subunits. This substrate specificity appears to be achieved by different modes of binding acyl-CoA: AccD5 strongly binds the CoA moiety but cannot accommodate long acyl chains, while AccD4 binds long acyl chains but does not interact strongly with CoA. Other propionyl-CoA carboxylase structures have been determined (Huang et al., 2010; Scheffen et al., 2021; Zhou et al., 2024), but the structure of the mycobacterial LCC complex has marked differences from these dedicated short-chain acyl-CoA carboxylases. Other propionyl-CoA carboxylases have a homo-oligomeric core of CT domain subunits and a 1:1:1 stoichiometry of BC:BCCP:CT domains. In contrast, the mycobacterial enzyme has a hetero-oligomeric core of CT domain subunits, more BC and BCCP domains than CT domains, and includes the unique AccE subunit as a tether. Despite the unusual architecture of the LCC complex, the AccD5 subunit appears to interact with propionyl-CoA and the BCCP domain of AccA3 in a manner comparable to other propionyl-CoA carboxylases. In all propionyl-CoA carboxylase structures determined to date, including the mycobacterial enzyme structures determined here under turnover conditions, biotin from the BCCP domain is resolved 8 to 9 Å from propionyl-CoA, which is too far for transfer of a carboxyl group to the substrate. A crystal structure of the *Streptomyces coelicolor* propionyl-CoA CT subunits with biotin soaked into the crystal showed the biotin ∼3.5 Å from the substrate, which was proposed to be the active state (Diacovich et al., 2004). It is possible that in the intact enzyme under turnover conditions, the enzyme conformation with biotin close to the acyl-CoA substrate is too short-lived to be observed by cryo-EM.

We found that one or two copies of a complex formed from MSMEG_0435 and MSMEG_0436 can bind to the LCC complex. MSMEG_0435 and MSMEG_0436 are homologous with *Bacillus subtilis* KipA/PxpC (Uniprot: Q7WY77) and KipI/PxpB (Uniprot: P60495), respectively (Fan et al., 2013, 2012). Orthologs of MSMEG_0435 and MSMEG_0436 are found in other mycobacteria, such as Rv0263c and Rv0264c in *M. tuberculosis* and ML2550 and ML2549 in *Mycobacterium leprae*. These genes and their homologs are annotated as allophanate hydrolase subunit 2 and 1, but it has already been noted that the proteins have no detectable sequence homology to the allophanate hydrolase component of urea amidolyase (Fan et al., 2012). Homologs of MSMEG_0435 and MSMEG_0436 have also been annotated as urea carboxylases (Petridis et al., 2015), 5-oxoprolinases (Niehaus et al., 2017), or inhibitors of a histidine kinase involved in sporulation signalling (Jacques et al., 2011; Wang et al., 1997). MSMEG_0435 has homology with urea carboxylases, but as expected from it missing key residues (Fan et al., 2012), we could not detect urea carboxylase activity *in vitro*. However, in a model of TB transmission, a recent screen of a genome-wide CRISPRi library found that Rv0263c was required for *M. tuberculosis* to survive rehydration in model alveolar lining fluid and subsequent uptake by alveolar macrophage-like cells (Singh et al., 2025). This observation suggests that MSMEG_0435 and MSMEG_0436, along with Rv0263c and Rv0264c in *M. tuberculosis*, may be involved in regulating the energetically costly process of synthesizing components of the cell wall, a process that requires regulation for successful infection by mycobacterial pathogens.

## Materials and methods

### Bacterial culture and isolation of membranes

The *M. smegmatis* strain SABM15, with sequence encoding a C-terminal 3×FLAG tag at the 3′ end of the gene for malate:quinone oxidoreductase (Di Trani et al., 2025), was cultured for 48-72 h at 30°C with 180 rpm shaking in 2.8 L Fernbach flasks containing 1 L of medium (Middlebrook 7H9 broth supplemented with 11 mM of sodium succinate and 0.05 % [v/v] tween-80). This medium allowed for growth of sufficient biomass while limiting contamination of the purified proteins with ribosomes. Cells were collected by centrifugation at 6,500 *g* for 20 min and resuspended at 15 mL/L in lysis buffer (50 mM 2-[N-morpholino]ethanesulfonic acid [MES] pH 6, 100 mM NaCl, 5 mM MgSO_4_, 5 mM benzamidine-HCl, 5 mM 6-aminocaproic acid) before lysis at >20 kPa with an Emulsiflex C3 homogenizer (Avestin). Cellular debris was removed by centrifugation at 39,000 *g* for 30 min. Membranes were collected by ultracentrifugation at 200,000 *g* for 60 min before being resuspended in 45 mL of buffer (20 mM MES pH 6, 100 mM NaCl, 5 mM CaCl_2_, 5 mM benzamidine-HCl, 5 mM 6-aminocaproic acid, 20% [v/v] glycerol) and frozen at -80°C.

### Protein purification

All protein purification was conducted at 4°C. Frozen membranes were thawed and filtered sequentially with 0.45 and 0.22 µm syringe filters (Millipore) before being loaded onto a plastic column containing 1 mL of Strep-Tactin XT Sepharose chromatography resin (Cytiva) equilibrated with binding buffer (20 mM MES pH 6, 100 mM NaCl, 2 mM CaCl_2_, 10 % [v/v] glycerol). The resin was washed with five column volumes of binding buffer and ten column volumes of sample buffer (20 mM MES pH 6, 100 mM NaCl, 2 mM CaCl_2_). Bound protein was eluted with 15 column volumes of 10 mM D-biotin in sample buffer. Eluted protein was concentrated with a 100 kDa cut-off concentrator (Millipore) and loaded onto a Superose 6 Increase 10/300 GL column (Cytiva) equilibrated with sample buffer. Fractions from the Superose 6 Increase column that contained the MCC and LCC complexes were pooled and concentrated with a 100 kDa cut-off concentrator and either used immediately to prepare cryo-EM grids or stored at 4°C for enzyme assays.

### Mass spectrometry sample preparation

The purified protein solution was diluted to 1 mg/mL in freshly prepared 50 mM ammonium bicarbonate. The sample was reduced by addition of 5 mM dithiothreitol at 56°C for 30 min and then alkylated with 50 mM iodoacetamide for 45 min in the dark. Following alkylation, 0.5 µg of sequencing grade modified trypsin (Promega) was added and the sample was incubated overnight at 37°C. The peptide solution was subsequently acidified with 5% formic acid before storing at -20°C. Frozen samples were sent to the Lunenfeld-Tanenbaum Network Biology Collaborative Centre Proteomics facility for LC-MS/MS mass spectrometry with a timsTOF Pro 2 mass spectrometer (Bruker), where the sample was processed and data-dependent analysis performed with their standard workflow (**Supplementary Data 1**).

### Activity assays

Enzyme activity was measured with a reduced nicotinamide adenine dinucleotide (NADH) oxidation-coupled ATP hydrolysis assay (Janiyani et al., 2001). Protein concentration was measured by bicinchoninic acid assay (BCA, Pierce) immediately before measurement of activity. To measure propionyl-CoA carboxylase activity, 8.5 µg/mL protein in sample buffer and 1 mM propionyl-CoA from a 8 mM stock in water were added to the assay mixture containing final concentrations of 50 mM MES pH 6.0, 100 mM NaCl, 3 mM MgCl_2_, 3.2 units of pyruvate kinase (Sigma), 8 units of lactate dehydrogenase (Sigma), and 200 µM NADH in a 96 well clear flat-bottom micro test plate (Sarstedt), with a final total volume of 160 µL/well. To measure arachidoyl-CoA carboxylase activity, 260 µg/mL protein in sample buffer and 500 µM arachidoyl-CoA from a 4 mM stock in water were added to the assay mixture containing final concentrations of 50 mM potassium phosphate pH 7.5, 100 mM NaCl, 3 mM MgCl_2_, 3.2 units of pyruvate kinase, 8 units of lactate dehydrogenase, and 200 µM NADH in a black 96 well half-area clear flat-bottom microplate (Corning), with a total volume of 80 µL/well. To detect urea carboxylase activity, 350 µg/mL protein in sample buffer and 50 mM urea from a 200 mM stock in water were added to the assay mixture containing final concentrations of 50 mM potassium phosphate pH 7.5, 100 mM NaCl, 3 mM MgCl_2_, 3.2 units of pyruvate kinase, 8 units of lactate dehydrogenase, and 200 µM NADH in a black 96 well half-area clear flat-bottom microplate (Corning), with a total volume of 80 µL/well. All assays were initiated by addition of 4 mM ATP, 2 mM phosphoenolpyruvic acid (PEP) (monopotassium salt, Sigma), and 50 mM NaHCO_3_ from a combined stock of 40 mM, 20 mM, and 500 mM, respectively. Reactions were monitored by measuring absorbance at 340 nm with either a SpectraMax M5e Multi-Mode Microplate Reader (Molecular Devices) or a BioTek Synergy Neo2 plate reader (Agilent) at 25°C.

### Cryo-EM specimen preparation

Holey gold foils with 2 µm holes in a square lattice were nanofabricated on 400 mesh copper– rhodium grids as described previously (Marr et al., 2014) and glow discharged for 15 s in air before use. Purified protein sample was concentrated to ∼5 mg/mL as measured by BCA assay. When imaged without substrates, 2 µL of the sample was applied on the grid in the environmentally controlled chamber of an EM GP2 Plunge Freezer (Leica Microsystems), maintained at 4°C and with a nominal ∼90% relative humidity. Grids were blotted for 1 s before plunge freezing in liquid ethane. For the sample imaged under turnover conditions with propionyl-CoA, 3 µL of propionyl-CoA was added to 9 µL of sample at a final concentration of 2 mM from an 8 mM stock dissolved in water and incubated for ∼15 min on ice. This mixture (1.8 µL) was applied to a holey gold grid in the grid freezing device and 0.2 µL solution of 20 mM ATP, 20 mM MgCl_2_, and 500 mM NaHCO_3_ that had been passed through a 0.22 µM filter was added and mixed on the grid. The grid was then blotted for 1 s before plunge freezing in liquid ethane. For the sample with both propionyl-CoA and arachidoyl-CoA, a stock solution of 8 mM propionyl-CoA and 4 mM arachidoyl-CoA was prepared in water. This solution (3 µL) was added to 9 µL of sample and incubated on ice for ∼15 min. The mixture (1.8 µL) was applied to a holey gold grid in the grid freezing device and 0.2 µL the solution with ATP, MgCl_2_, and NaHCO_3_ was added, mixed, blotted for 1 s, and the grid was plunge frozen in liquid ethane.

### Cryo-EM imaging

Both specimen screening and high-resolution data collection were automated with the EPU software (ThermoFisher Scientific). All samples were screened with a Glacios 2 electron microscope operating at 200 kV and equipped with a Falcon 4i camera (ThermoFisher Scientific). Images were collected in electron event representation (EER) mode (Guo et al., 2020) at 92,000× nominal magnification, corresponding to a calibrated pixel size of 1.5 Å, with a total exposure of ∼40 e^−^/Å^2^. For high-resolution data collection of all samples, 8,000 movies were collected with a Titan Krios G3 electron microscope operating at 300 kV and equipped with a Falcon4i camera and SelectrisX energy filter (ThermoFisher Scientific), using a slit width of 10 eV. The nominal magnification was 130,000×, corresponding to a calibrated pixel size of 0.93 Å. For the substrate-free and propionyl-CoA-bound datasets, a total exposure of ∼40 e^−^/Å^2^ was used. For the dataset with both propionyl-CoA and arachidoyl-CoA, a total exposure of ∼50 e^−^/Å^2^ was used.

### Cryo-EM image analysis

Data quality during automated collection was monitored with cryoSPARC Live. All image analysis was done with cryoSPARC v5.0.0-privatebeta (Punjani et al., 2017). Exposure fractions were aligned (Rubinstein and Brubaker, 2015), and contrast transfer function (CTF) parameters estimated in patches. All refinements were done without imposing symmetry except where stated.

For the LCC map without substrate (**Fig. S3**), initial particle selection was performed with the blob picker from a subset of 990 aligned movies. The particle images were extracted with a box size of 480×480 pixels, Fourier cropped to 120×120 pixels (3.72 Å/pixel), and subjected to two-dimensional (2D) classification. Initial maps from particle images in selected 2D classes were calculated by *ab initio* reconstruction and cleaned by heterogeneous refinement. The particle images that contributed to the best map were then subjected to 2D classification to generate templates for template picker on denoised micrographs. Following use of template picker on the full dataset, 2,419,931 particle images were extracted with a box size of 480×480 pixels and Fourier cropped to 120×120 pixels (3.72 Å/pixel). This dataset was cleaned with one round of 2D classification and initial maps were calculated by *ab initio* reconstruction and subsequent heterogeneous refinement. 778,798 particle images corresponding to the best 3D map were re-extracted with a box size of 480×480 pixels (0.93 Å/pixel). This set of particle images was cleaned with multiple iterations of *ab initio* reconstruction and heterogeneous refinement before a high-resolution map was calculated with non-uniform refinement (Punjani et al., 2020) from 131,774 particle images. Individual particle CTF parameters were estimated, and the particle images were subject to an additional round of non-uniform refinement before 3D classification without a focused mask, which was used to separate particle images where the full AccA3_8_AccD4_2_AccD5_4_AccE_2_ complex could be resolved. These 11,174 particle images were used to calculate the final high-resolution map by non-uniform refinement. The region of the map with the flexible AccA3 subunits were subjected to local refinement.

For the LCC map with propionyl-CoA bound, a workflow like the workflow for the substrate-free map was followed (**Fig. S7**). Initial particle coordinates were generated from blob picker from a subset of 2,000 micrographs. Templates for template picking on denoised micrographs were generated in a similar manner but were extracted with a box that was Fourier cropped to 240×240 pixels (1.86 Å/pixel) instead. Following use of template picker, 2,018,368 particle images were extracted with a box size of 480×480 pixels and Fourier cropped to 240×240 pixels (1.86 Å/pixel). The dataset was cleaned with one round of 2D classification and initial maps were calculated by *ab initio* reconstruction and heterogeneous refinement. 693,566 particle images from the best map were re-extracted with a box size of 480×480 pixels (0.93 Å/pixel).

This dataset was cleaned with multiple iterations of *ab initio* reconstruction and heterogeneous refinement. A high-resolution map was calculated from 148,465 particle images and refined with non-uniform refinement. Individual particle CTF parameters were estimated and the particle images were subjected to an additional round of non-uniform refinement. For the map with the BCCP domains in the “down” conformation, particle images where the full AccA3_8_AccD4_2_AccD5_4_AccE_2_ complex could be resolved were separated by 3D classification without a focused mask. These 46,806 particle images were used to calculate a high-resolution map with non-uniform refinement, and the map regions corresponding to the flexible AccA3 subunits were locally refined. For maps with the BCCP domains in the “up” conformation, the 148,465 particle images were first subjected to local refinement with a mask that included the two AccA3 subunits on one side of the AccD4_2_AccD5_4_ hexamer. 3D classification with a mask that included the region where the BCCP domains interact with the BC domains allowed for separation of particle images with clear BCCP-BC interactions. A 3D map was calculated with non-uniform refinement and the final high-resolution map with two BCCP domains in the “up” conformation was calculated from 20,330 particle images using local refinement with a mask that included the two AccA3 subunits on one side of the AccD4_2_AccD5_4_ hexamer. Similarly, two high-resolution locally-refined maps where each of the two BCCP domain was in the “up” conformation were calculated from 27,091 and 25,933 particle images, respectively.

For the LCC map with both arachidoyl-CoA and propionyl-CoA bound, a similar workflow was again followed (**Fig. S10**). After use of template picker on denoised micrographs, 2,514,602 particle images were extracted with a box size of 480×480 pixels and Fourier cropped to 128×128 pixels (3.49 Å/pixel). This dataset was cleaned by one round of 2D classification and initial maps were calculated *ab initio* and subjected to heterogeneous refinement. The 736,301 particle images that contributed to this map were re-extracted with a box size of 480×480 pixels (0.93 Å/pixel). This particle image set was cleaned with several iterations of *ab initio* reconstruction and heterogeneous refinement and a high-resolution map was calculated with non-uniform refinement from 542,340 particle images. Individual particle CTF parameters were estimated and the map was subject to another round of non-uniform refinement. Overrepresented views were removed by 3D classification without a focused mask followed by non-uniform refinement of the map. Particle images that produced a map with the complex of MSMEG_0435-MSMEG_0436 bound were separated by 3D classification with a mask that included the AccD4_2_AccD5_4_ hexamer core. The remaining particle images were again subjected to 3D classification without a focused mask to separate particle images containing the full AccA3_8_AccD4_2_AccD5_4_AccE_2_ complex. These 124,481 particle images were used to calculate the final high-resolution map with non-uniform refinement and the flexible AccA3 region of the map was subjected to local refinement. The particle image dataset that showed MSMEG_0435-MSMEG_0436 complexes were cleaned by iterations of 3D classification and non-uniform refinement to yield two maps of the LCC complex. One of these maps had one MSMEG_0435-MSMEG_0436 complex bound and was calculated from 56,830 particle images. The other had two MSMEG_0435-MSMEG_0436 complexes bound and was calculated from 10,016 particle images.

A map of AccA1_6_AccD1_6_ MCC complex could be determined from each of the datasets, but the dataset with both propionyl-CoA and arachidoyl-CoA gave the highest resolution map (**Fig. S10**). Particle images were selected from the dataset obtained following use of the template picker with templates corresponding to 2D class averages showing the LCC complex. These particle images were extracted and Fourier cropped to a box size of 128×128 pixels (3.49 Å/pixel). Initial maps were generated by *ab initio* reconstruction, cleaned with heterogeneous refinement, and 108,486 particle images were re-extracted with a box size of 480×480 pixels (0.93 Å/pixel). Particle images were cleaned with one round of *ab initio* reconstruction and heterogeneous refinement and a high-resolution map with D3 symmetry was calculated with non-uniform refinement from 87,410 particle images. Overrepresented views were removed with 3D classification, and the final high-resolution map was calculated with non-uniform refinement from 30,234 particle images with D3 symmetry imposed.

### Model building and refinement

For the map of the LCC complex without substrate and MCC complex, initial atomic models were generated by ModelAngelo (Jamali et al., 2024) with amino acid sequences provided. All subsequent LCC maps used this model as the starting model. For the map of the LCC complex with MSMEG_0435 and MSMEG_0436 bound, a crystal structure of the complex of MSMEG_0435 and MSMEG_0436 (PDB: 3MML) was used as the starting model for the subcomplex. Large gaps in the model were filled with segments of protomers obtained from the AlphaFoldDB protein structure database (Jumper et al., 2021; Varadi et al., 2022) and fit as rigid bodies into the map with UCSF ChimeraX (Goddard et al., 2018; Meng et al., 2023). A 3D model of arachidoyl-CoA was generated with *phenix.elbow* from the SMILES (Simplified Molecular Input Line Entry System) string. Combining model fragments, optimisation of model-to-map fit and molecular dihedral angles, filling of small gaps in the starting model, and fitting of ligands was done with Coot v0.9.6 (Emsley and Cowtan, 2004) and ISOLDE v1.8 (Croll, 2018). Model parameters were refined with PHENIX v1.19.2 (Liebschner et al., 2019). All figures were rendered with UCSF ChimeraX.

## Author contributions

YL and JLR designed the experiments. YL performed all experiments and JLR supervised the research. YL and JLR wrote the manuscript and prepared the figures.

## Data, Materials, and Software Availability

The electron cryomicroscopy maps and associated models described in this article have been deposited in the Electron Microscopy Data Bank (EMDB) with accession numbers EMD-73568 (MCC complex, nonuniform refinement), EMD- 73569 (LCC complex without substrates, non-uniform refinement), EMD-73570 (LCC complex without substrates, local refinement of upper two AccA3 BC domains), EMD-73571 (LCC complex without substrates, local refinement of upper one AccA3 BC domain), EMD-73572 (LCC complex without substrates, local refinement of lower two AccA3 BC domains), EMD- 73573 (LCC complex without substrates, local refinement of lower one AccA3 BC domain), EMD-73575 (LCC complex without substrates, composite map), EMD-73579 (LCC complex with propionyl-CoA, non-uniform refinement), EMD-73580 (LCC complex with propionyl-CoA, local refinement of upper two AccA3 BC domains), EMD-73582 (LCC complex with propionyl-CoA, local refinement of upper one AccA3 BC domain), EMD-73583 (LCC complex with propionyl-CoA, local refinement of lower two AccA3 BC domains), EMD-73584 (LCC complex with propionyl-CoA, local refinement of lower one AccA3 BC domain), EMD-73585 (LCC complex with propionyl-CoA, composite map), EMD-73586 (LCC complex, left BCCP domain up), EMD-73587 (LCC complex, right BCCP domain up), EMD-73588 (LCC complex both BCCP domains up), EMD-73590 (LCC complex with arachidoyl-CoA/propionyl-CoA, non-uniform refinement), EMD-73591 (LCC complex with arachidoyl-CoA/propionyl-CoA, local refinement of upper two AccA3 BC domains), EMD-73592 (LCC complex with arachidoyl-CoA/propionyl-CoA, local refinement of upper one AccA3 BC domain), EMD-73593 (LCC complex with arachidoyl-CoA/propionyl-CoA, local refinement of lower two AccA3 BC domains), EMD-73594 (LCC complex with arachidoyl-CoA/propionyl-CoA, local refinement of lower one AccA3 BC domain), EMD-73595 (LCC complex with arachidoyl-CoA/propionyl-CoA, composite map), and EMD-73596 (LCC complex with MSMEG_0435-MSMEG_0436 bound); and the Protein Data Bank with accession codes PDB_00009YX0 (MCC complex), PDB_00009YX1 (LCC complex without substrates), PDB_00009YX2 (LCC complex with propionyl-CoA), PDB_00009YX4 (LCC complex with arachidoyl-CoA/propionyl-CoA), and PDB_00009YX5 (LCC complex with MSMEG_0435-MSMEG_0436 bound).

## Acknowledgements

We thank Prof. Carl Nathan for helpful discussions on the CRISPRi screen that detected proteins necessary for TB transmission. We thank Samir Benlekbir for assistance with cryo-EM data collection and Cassandra Wong for mass spectrometry data collection and analysis. YL was supported by a Canada Graduate Scholarship - Doctoral program award from the Canadian Institutes of Health Research (CIHR) and JLR was supported by the Canada Research Chairs program. Research was funded by CIHR Project Grant PJT191893 (JLR). Cryo-EM data were collected at the Toronto High-Resolution High-Throughput cryo-EM Facility at The Hospital for Sick Children. Mass spectrometry data were collected at the Network Biology Collaborative Center at the Lunenfeld-Tanenbaum Research Institute. Some of the assays performed used infrastructure from the Structural and Biophysical Core Facility at The Hospital for Sick Children. These core facilities were supported by the Canada Foundation for Innovation and the Ontario Government.

---

**Supplementary Table 1.** Cryo-EM data collection and map refinement for the dataset with ATP, bicarbonate, arachidoyl-CoA, and propionyl-CoA.

|  | Collection details |  |  |  |  |  |  |
| --- | --- | --- | --- | --- | --- | --- | --- |
| Magnification | 130,000× |  |  |  |  |  |  |
| Voltage (kV) | 300 |  |  |  |  |  |  |
| Energy filter slit width (eV) | 10 |  |  |  |  |  |  |
| Micrographs collected | 8000 |  |  |  |  |  |  |
| Electron exposure (e<sup>-</sup>/Å<sup>2</sup>) | 50 |  |  |  |  |  |  |
| Nominal defocus range (μm) | 1.1–2.0 |  |  |  |  |  |  |
| Raw pixel size (Å) | 0.93 |  |  |  |  |  |  |
|  | Reconstruction details |  |  |  |  |  |  |
|  | Non-uniform refinement (EMD-73590) | Two AccA3s, upper (EMD-73591) | One AccA3, upper (EMD-73592) | Two AccA3s, lower (EMD-73593) | One AccA3, lower (EMD-73594) | LCC with two MSMEG_0435/MSMEG_0436 complexes (EMD-73596) | MCC (EMD-73568) |
| Symmetry imposed | C1 |  |  |  |  | C1 | D3 |
| Initial particle images extracted (no.) | 2,514,632 |  |  |  |  | 2,514,632 (same initial extraction as LCC) | 2,514,632 (same initial extraction as LCC) |
| Initial particle images for 3D processing (no.) | 535,937 |  |  |  |  | 535,937 (same extraction as core LCC) | 109,170 |
| Final particle images (no.) | 124,481 |  |  |  |  | 10,016 | 30,234 |
|  | Refinement details |  |  |  |  |  |  |
| Map resolution (Å) | 2.0 | 2.5 | 2.6 | 2.4 | 2.5 | 2.9 | 2.0 |
| FSC threshold | 0.143 | 0.143 | 0.143 | 0.143 | 0.143 | 0.143 | 0.143 |
| Total map resolution range (Å) | 2.1–34.0 | 2.2–44.4 | 2.3–44.2 | 2.2–44.4 | 2.3–44.1 | 2.5–44.6 | 2.1–31.9 |
| FSC mask map resolution range (Å) | 2.1–2.2 | 2.2–2.8 | 2.3–2.9 | 2.2–2.7 | 2.2–2.8 | 2.5–3.2 | 2.1–2.2 |
| cFAR | 0.65 | 0.6 | 0.67 | 0.61 | 0.62 | 0.51 | 0.75 |
| Map sharpening B-factor | 29.8 | 43.8 | 59.8 | 41.8 | 53.9 | 15.1 | 33.8 |

**Supplementary Table 2.** Cryo-EM data collection and map refinement for the dataset without substrate.

|  | Collection details |  |  |  |  |
| --- | --- | --- | --- | --- | --- |
| Magnification | 130,000× |  |  |  |  |
| Voltage (kV) | 300 |  |  |  |  |
| Energy filter slit width (eV) | 10 |  |  |  |  |
| Micrographs collected | 8000 |  |  |  |  |
| Electron exposure (e<sup>-</sup>/Å<sup>2</sup>) | 40 |  |  |  |  |
| Nominal defocus range (μm) | 1.1–2.0 |  |  |  |  |
| Raw pixel size (Å) | 0.93 |  |  |  |  |
|  | Reconstruction details |  |  |  |  |
| Symmetry imposed | C1 |  |  |  |  |
| Initial particle images extracted (no.) | 2,419,931 |  |  |  |  |
| Initial particle images for 3D processing (no.) | 131,774 |  |  |  |  |
| Final particle images (no.) | 11,174 |  |  |  |  |
|  | Refinement details |  |  |  |  |
|  | Non-uniform refinement (EMD-73569) | Two AccA3s, upper (EMD-73570) | One AccA3, upper (EMD-73571) | Two AccA3s, lower (EMD-73572) | One AccA3, lower (EMD-73573) |
| Map resolution (Å) | 2.9 | 3.3 | 3.4 | 3.4 | 4.0 |
| FSC threshold | 0.143 | 0.143 | 0.143 | 0.143 | 0.143 |
| Total map resolution range (Å) | 2.6–49.5 | 2.0–59.3 | 3.1–59.1 | 3.1–59.4 | 3.5–63.7 |
| FSC mask map resolution range (Å) | 2.6–3.3 | 2.8–4.0 | 3.1–4.0 | 3.1–4.1 | 3.5–7.2 |
| cFAR | 0.42 | 0.36 | 0.3 | 0.31 | 0.23 |
| Map sharpening B-factor | 18.2 | 21.4 | 33.5 | 20.7 | 55.0 |

**Supplementary Table 3.** Cryo-EM data collection and map refinement for the dataset with ATP, bicarbonate, and propionyl-CoA.

|  | Collection details |  |  |  |  |  |  |  |
| --- | --- | --- | --- | --- | --- | --- | --- | --- |
| Magnification | 130,000× |  |  |  |  |  |  |  |
| Voltage (kV) | 300 |  |  |  |  |  |  |  |
| Energy filter slit width (eV) | 10 |  |  |  |  |  |  |  |
| Micrographs collected | 8000 |  |  |  |  |  |  |  |
| Electron exposure (e<sup>-</sup>/Å<sup>2</sup>) | 40 |  |  |  |  |  |  |  |
| Nominal defocus range (μm) | 1.1–2.0 |  |  |  |  |  |  |  |
| Raw pixel size (Å) | 0.93 |  |  |  |  |  |  |  |
|  | Reconstruction details |  |  |  |  |  |  |  |
| Symmetry imposed | C1 |  |  |  |  |  |  |  |
| Initial particle images extracted (no.) | 2,018,368 |  |  |  |  |  |  |  |
| Initial particle images for 3D processing (no.) | 148,465 |  |  |  |  |  |  |  |
|  | Non-uniform refinement (EMD-73579) | Two AccA3s, upper (EMD-73580) | One AccA3, upper (EMD-73582) | Two AccA3s, lower (EMD-73583) | One AccA3, lower (EMD-73584) | Left BCCP up (EMD-73586) | Right BCCP up (EMD-73587) | Both BCCPs up (EMD-73589) |
| Final particle images (no.) | 46,806 |  |  |  |  | 25,933 | 27,091 | 20,330 |
|  | Refinement details |  |  |  |  |  |  |  |
| Map resolution (Å) | 2.3 | 2.8 | 3.0 | 2.8 | 2.9 | 3.3 | 3.2 | 3.4 |
| FSC threshold | 0.143 | 0.143 | 0.143 | 0.143 | 0.143 | 0.143 | 0.143 | 0.143 |
| Total map resolution range (Å) | 2.1–39.4 | 2.6–49.3 | 2.6–49.3 | 2.5–49.4 | 2.7–49.4 | 2.8–54.0 | 3.0–58.4 | 2.0–58.6 |
| FSC mask map resolution range (Å) | 2.1–2.7 | 2.6–3.2 | 2.6–3.6 | 2.5–3.2 | 2.7–3.3 | 2.8–3.8 | 3.0–3.7 | 3.0–4.1 |
| cFAR | 0.51 | 0.53 | 0.46 | 0.51 | 0.49 | 0.36 | 0.50 | 0.37 |
| Map sharpening B-factor | 29.9 | 41.7 | 60.8 | 41.4 | 54.1 | 46.0 | 48.6 | 48.3 |

**Supplementary Table 4:** Atomic model refinement and validation.

| Name (PDB accession, EMD accession) | MCC (9YX0, EMD-73568) | LCC without substrate (9YX1, EMD-73575) | Propionyl-CoA-bound LCC (9YX2, EMD-73585) | Arachidoyl-CoA/Propionyl-CoA-bound LCC (9YX4, EMD-73595) | LCC with MSMEG_0435-MSMEG_0436 (9YX5, EMD-73596) |
| --- | --- | --- | --- | --- | --- |
| **Model building and refinement** |  |  |  |  |  |
| Initial model used | ModelAngelo | ModelAngelo | Built model of LCC without substrate and AlphaFoldDB | Built model of LCC with propionyl-CoA | Built model of LCC with propionyl-CoA and PDB: 3MML |
| Model resolution (Å) | 2.21 | 3.17 | 2.65 | 2.04 | 3.17 |
| FSC threshold | 0.5 | 0.5 | 0.5 | 0.5 | 0.5 |
| CC (volume/mask) | 0.86/0.86 | 0.84/0.84 | 0.87/0.87 | 0.88/0.88 | 0.80/0.82 |
| Protein B-factors (Å<sup>2</sup>) | 78.83 | 86.79 | 67.46 | 58.61 | 114.68 |
| Ligand B-factors (Å<sup>2</sup>) | 68.62 | 75.89 | 75.38 | 62.34 | 63.78 |
| **Model composition** |  |  |  |  |  |
| Nonhydrogen atoms | 47,381 | 48,731 | 54,893 | 56,623 | 56,321 |
| Protein residues | 6588 | 7040 | 7657 | 7663 | 8786 |
| Ligands | BTI: 6 | BCT: 2; BTI: 6 | MG: 8; ATP: 8; BCT: 8; 1VU: 4; BTI: 6 | MG: 8; A1CZD: 2; ATP: 8; BCT: 8; 1VU: 4; BTI: 6 | MG: 2; A1CZD: 2; ATP: 2; BCT: 2; 1VU: 4; BTI: 5 |
| **RMS deviations** |  |  |  |  |  |
| Bond lengths (Å) | 0.006 | 0.004 | 0.010 | 0.009 | 0.006 |
| Bond angles (°) | 0.771 | 0.826 | 0.774 | 0.889 | 0.908 |
| **Validation** |  |  |  |  |  |
| MolProbity score | 0.88 | 1.50 | 1.14 | 1.21 | 1.19 |
| Clashscore | 0.93 | 3.93 | 3.20 | 2.22 | 1.65 |
| Poor rotamers (%) | 0.76 | 1.02 | 0.08 | 0.73 | 0.63 |
| CaBLAM outliers (%) | 1.14 | 1.44 | 1.69 | 1.73 | 1.89 |
| EMRinger Score | 6.37 | 4.73 | 5.54 | 6.23 | 4.74 |
| **Ramachandran plot** |  |  |  |  |  |
| Favoured (%) | 97.53 | 95.53 | 97.87 | 96.62 | 95.98 |
| Allowed (%) | 2.47 | 4.18 | 2.10 | 3.35 | 4.02 |
| Disallowed (%) | 0.00 | 0.29 | 0.03 | 0.03 | 0.00 |

**Supplementary Data 1.** Mass spectrometry of purified biotinylated proteins.

**Supplementary Movie 1.** Atomic model for BCCP translocation from the “down” conformation to the “up” conformation.

**Supplementary Figure 1.**
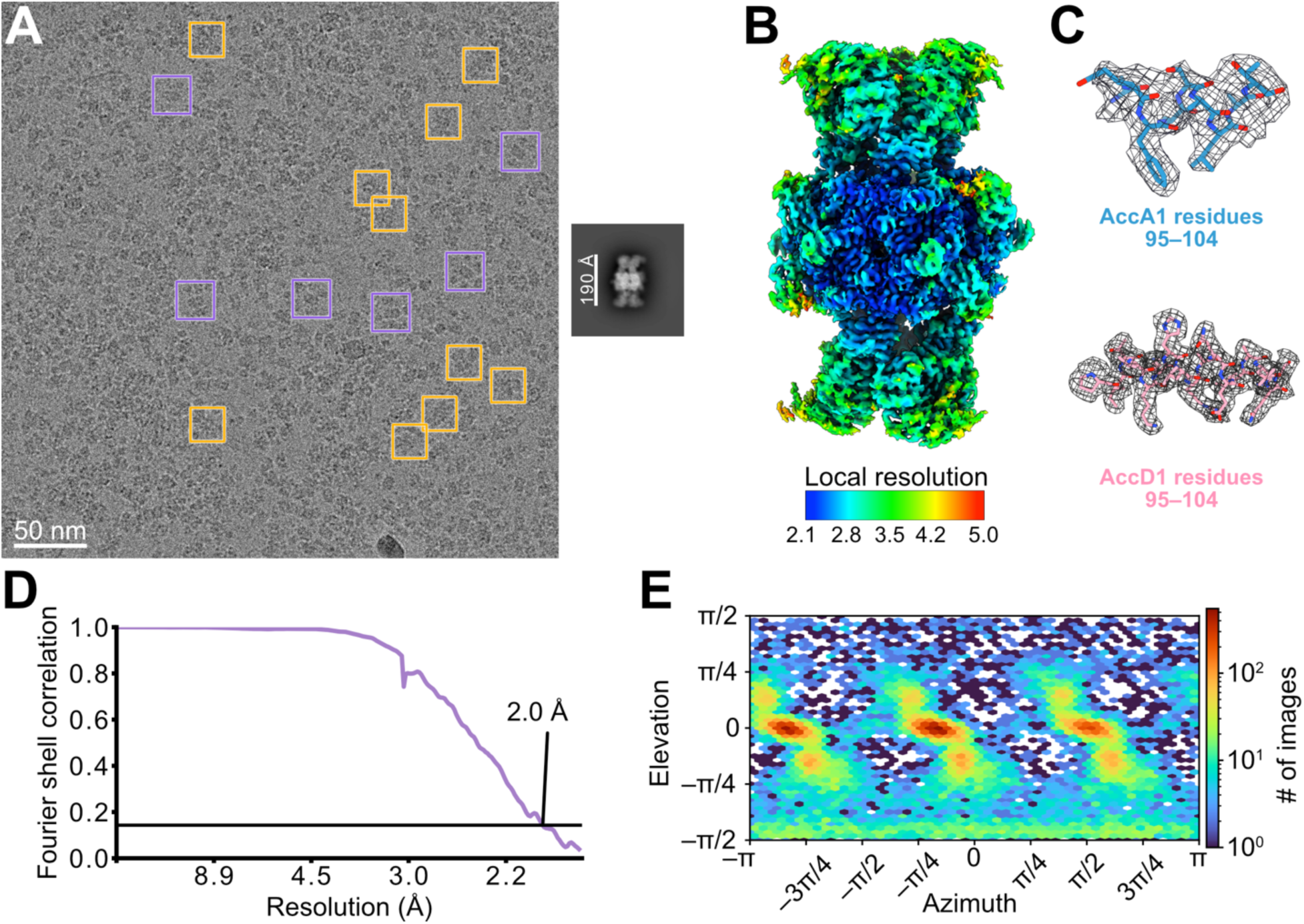
Cryo-EM map validation for the 3-methylcrotonyl-CoA carboxylase (MCC) complex. **A,** Example micrograph (*left*). Selected long-chain acyl-CoA carboxylase (LCC) complexes (*purple boxes*) and MCC complexes (*yellow boxes*) are indicated. 2D class average image (*right*) showing the MCC complex. **B,** Local resolution map for the MCC complex. **C,** Example model-in-map fit for selected regions of the MCC complex. **D,** Fourier shell correlation curve following a gold-standard refinement and correction for the effects of masking. **E,** Orientation distribution plot for particle images.

**Supplementary Figure 2.**
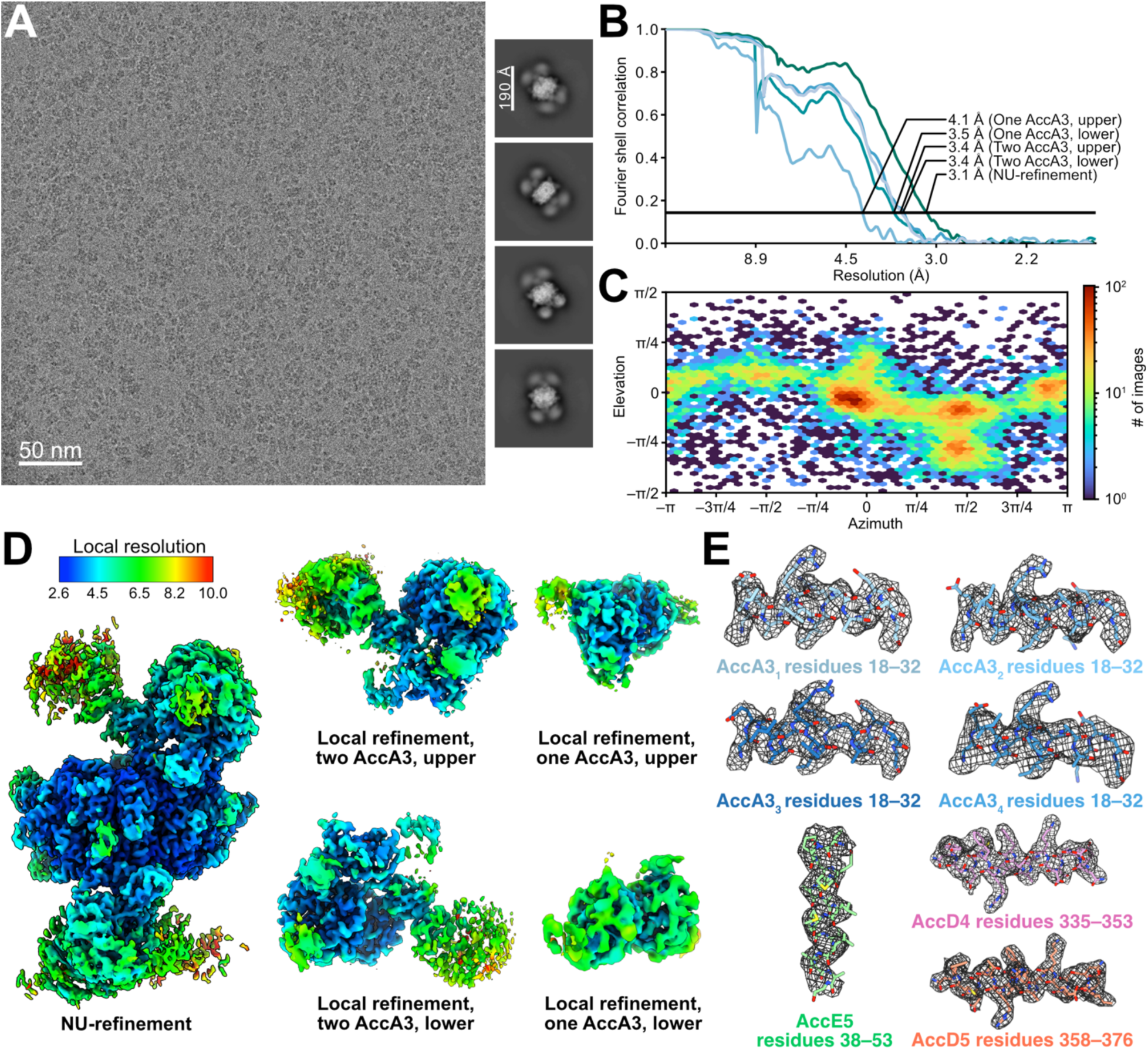
Cryo-EM map validation for the long-chain acyl-CoA carboxylase (LCC) complex without substrates. **A,** Example micrograph (*left*), and 2D class average images (*right*) showing the LCC complex. **B,** Fourier shell correlation curves following a gold-standard refinement and correction for the effects of masking for the overall non-uniform (NU) refinement of the map and local refinement of AccA3 BC domains. **C,** Orientation distribution plot for particle images. **D,** Local resolution maps for the different regions of the LCC complex. **E,** Example model-in-map fit for selected regions of the LCC complex.

**Supplementary Figure 3.**
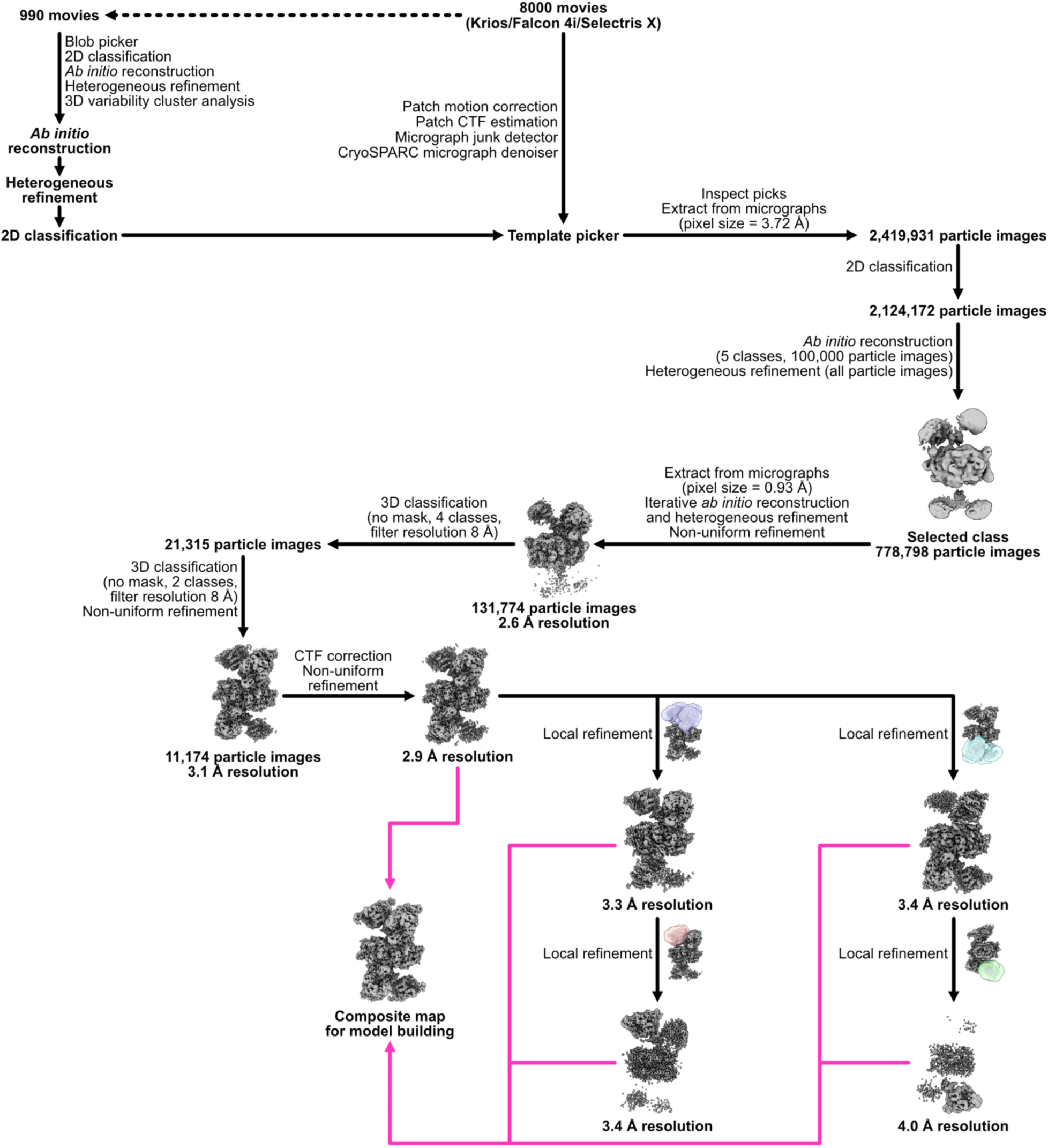
Workflow for calculating maps of the long-chain acyl-CoA carboxylase (LCC) complex without substrates.

**Supplementary Figure 4.**
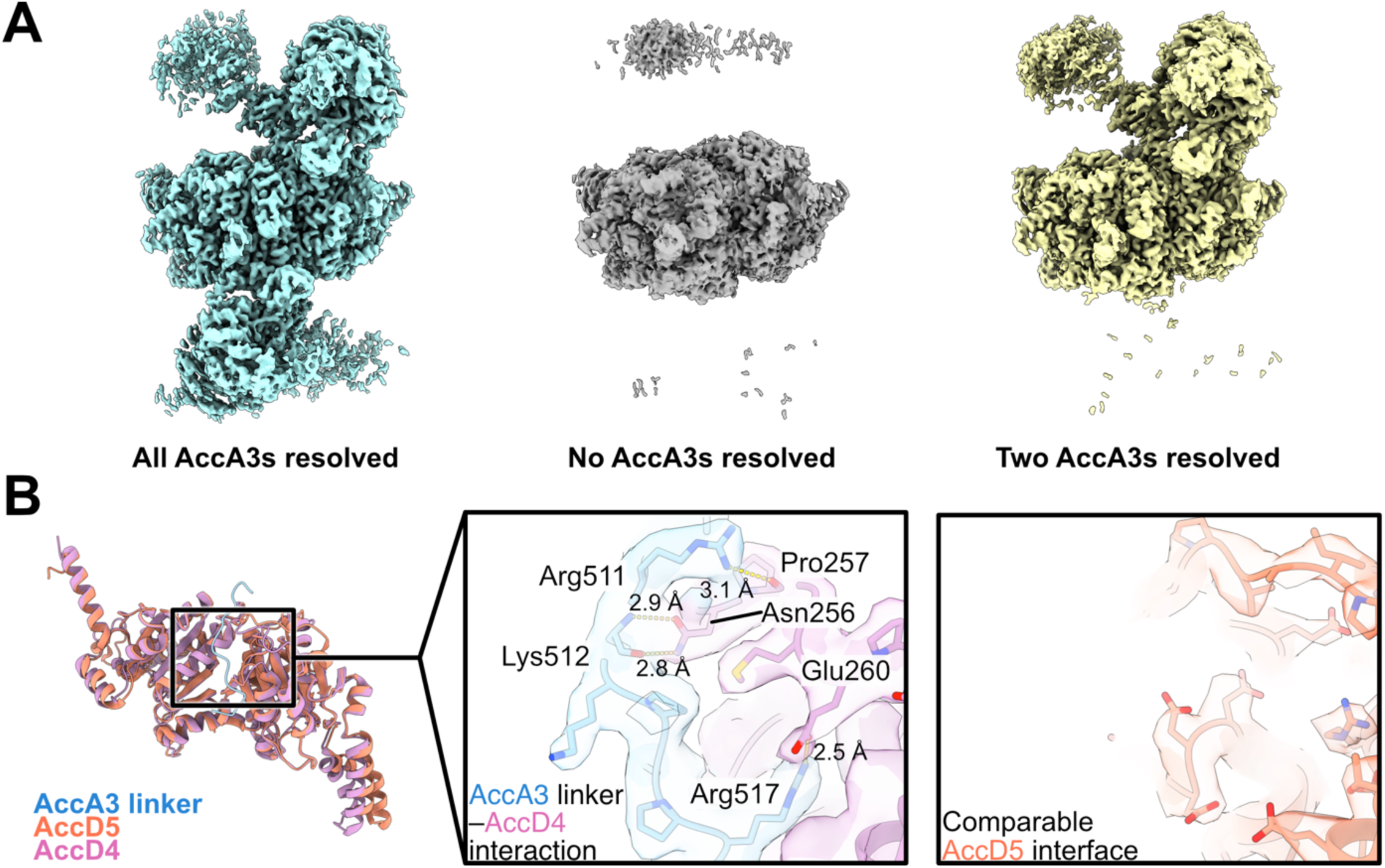
Stoichiometry and interactions of the AccA3 subunits in the LCC complex. **A,** 3D classes of the LCC complex with six (*left*), no (*middle*), and two (*right*) AccA3 subunits resolved. **B,** Interaction of the AccA3 subunit linker (residues 508 to 517) with AccD5 (*middle*) compared to AccD4 (*right*).

**Supplementary Figure 5.**
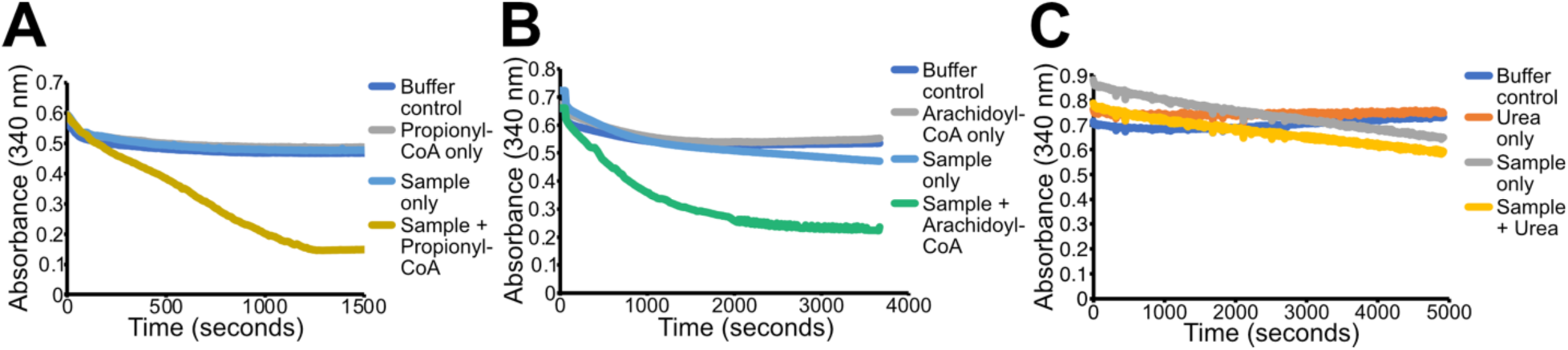
Replicate carboxylation assays. **A,** Propionyl-CoA carboxylation assay with the protein sample shows activity. **B,** Arachidoyl-CoA carboxylation assay with the protein sample shows activity. **C,** Urea carboxylase assay with the protein sample shows no activity.

**Supplementary Figure 6.**
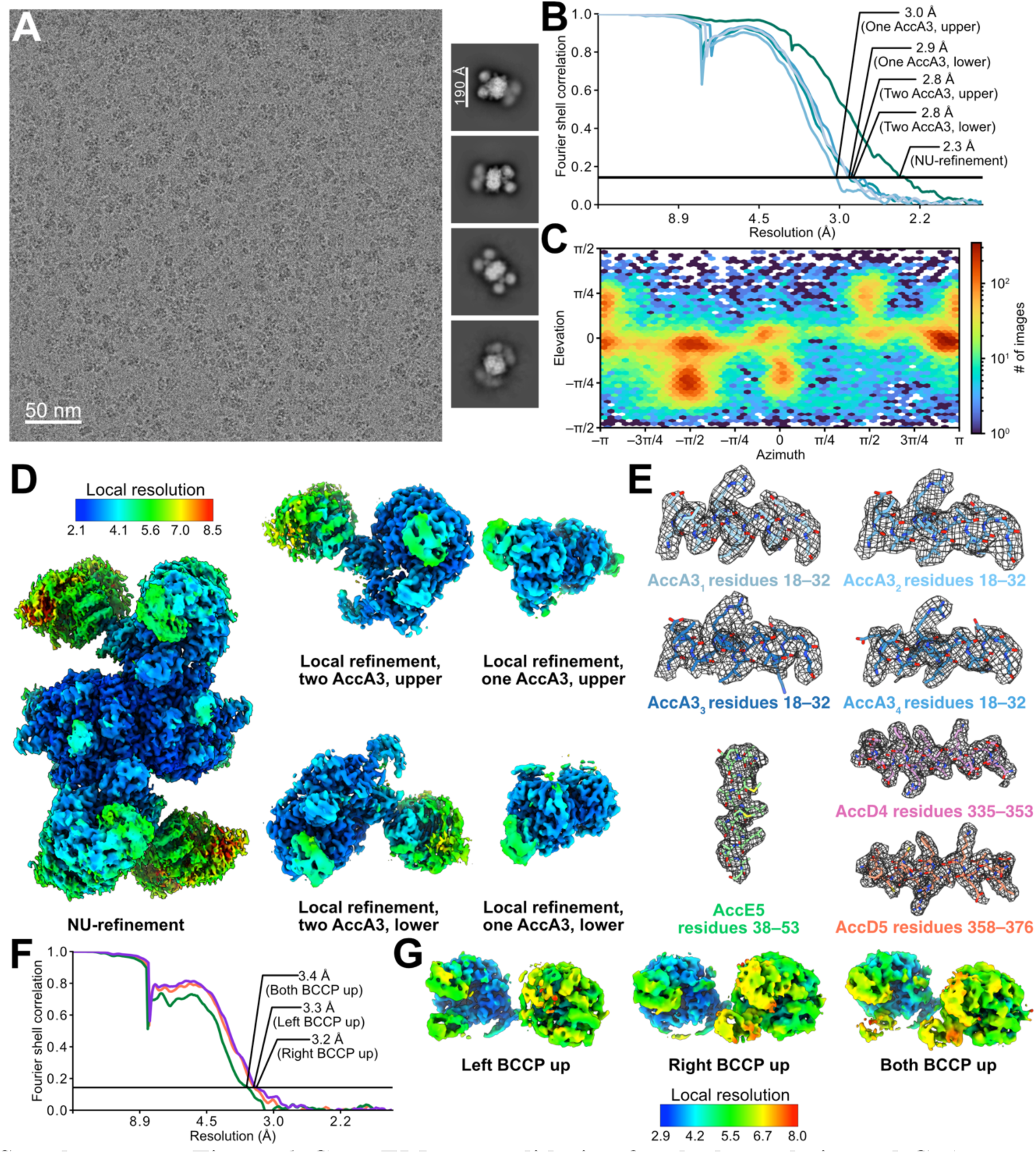
Cryo-EM map validation for the long-chain acyl-CoA carboxylase (LCC) complex with ATP, bicarbonate, and propionyl-CoA. **A,** Example micrograph (*left*), and 2D class average images (*right*) showing the LCC complex in the presence of ATP, bicarbonate, and propionyl-CoA. **B,** Fourier shell correlation curves following a gold-standard refinement and correction for the effects of masking for the overall non-uniform (NU) refinement, and with correction for the effects of masking for local refinement of AccA3 BC domains. **C,** Orientation distribution plot for particle images. **D,** Local resolution maps for the different regions of the LCC complex with ATP, bicarbonate, and propionyl-CoA. **E,** Example model-in-map fit for selected regions of the LCC complex with ATP, bicarbonate, and propionyl-CoA. **F,** Fourier shell correlation curves with correction for the effects of masking for local refinement of the BCCP domains in the “up” conformation. **G,** Local resolution maps for the BCCP domains in the “up” conformation.

**Supplementary Figure 7.**
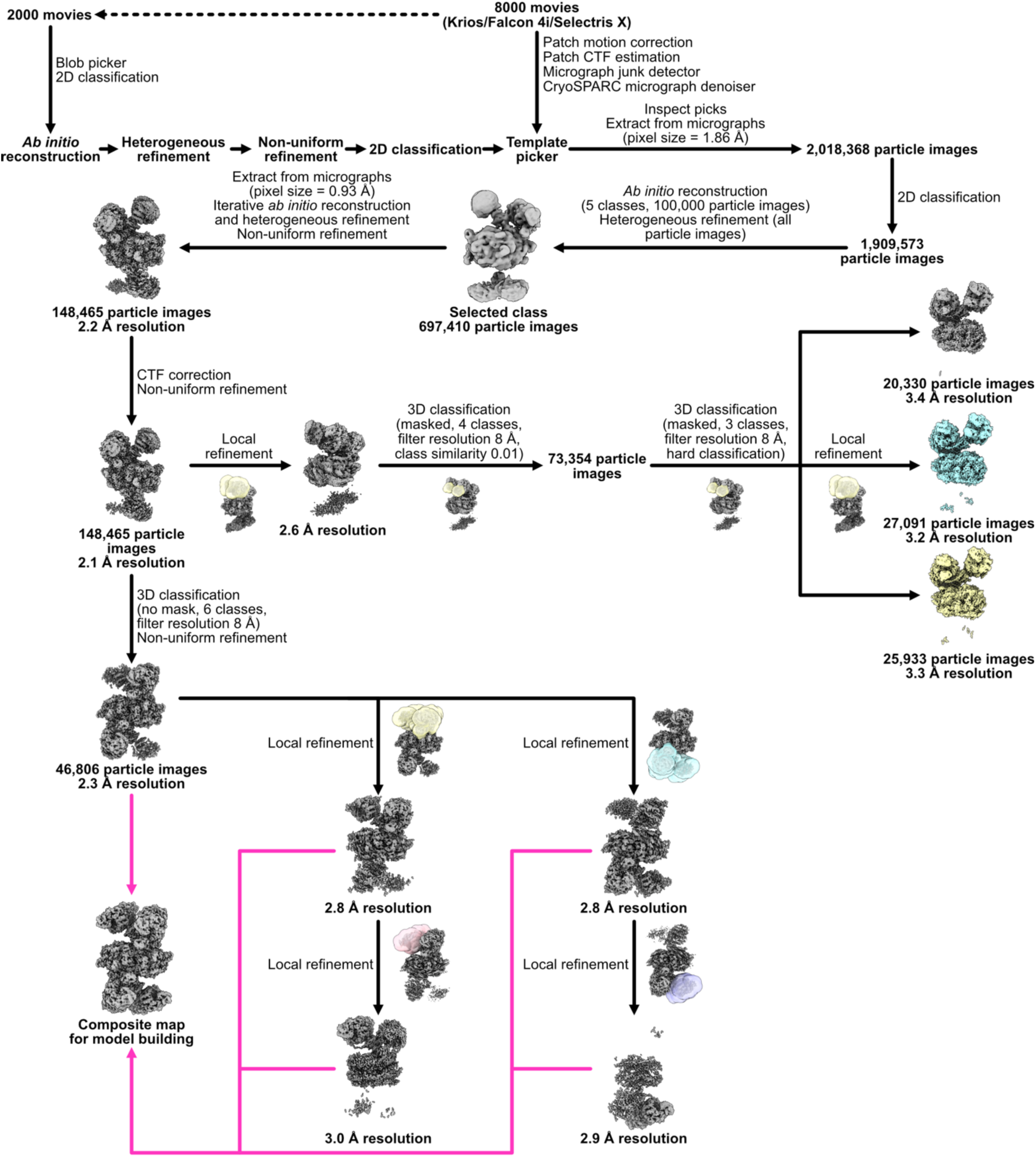
Workflow for calculating maps of the long-chain acyl-CoA carboxylase (LCC) complex with ATP, bicarbonate, and propionyl-CoA.

**Supplementary Figure 8.**
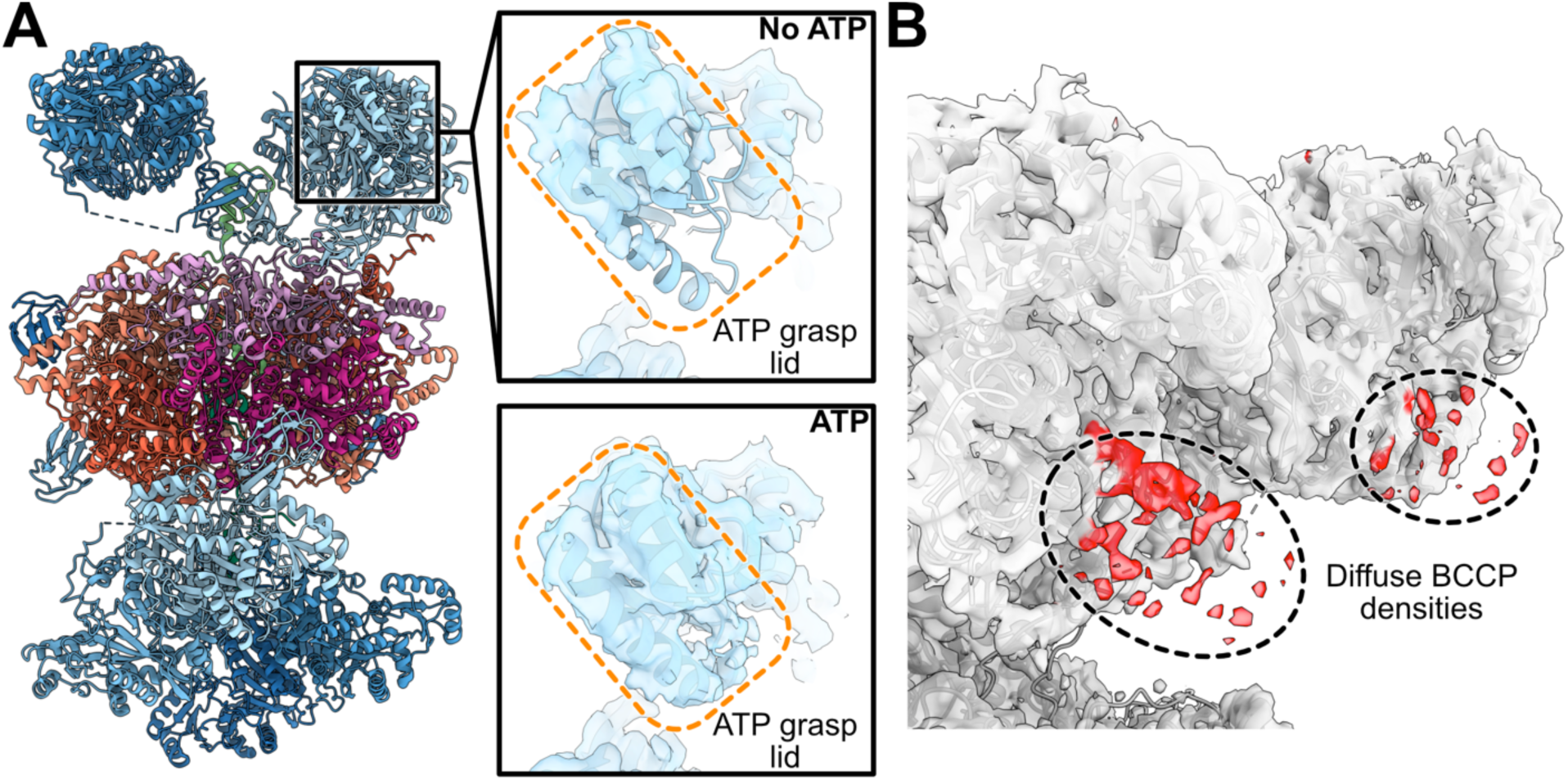
Conformational changes in the LCC complex in the presence of ATP, bicarbonate, and propionyl-CoA. **A,** The lid of the ATP grasp motif in the BC domain of AccA3 becomes better ordered in the presence of ATP. **B,** Diffuse density from the BCCP domains (*red*) is seen adjacent to the BC domains near the AccD5 subunits when the sample is frozen in the presence of substrate.

**Supplementary Figure 9.**
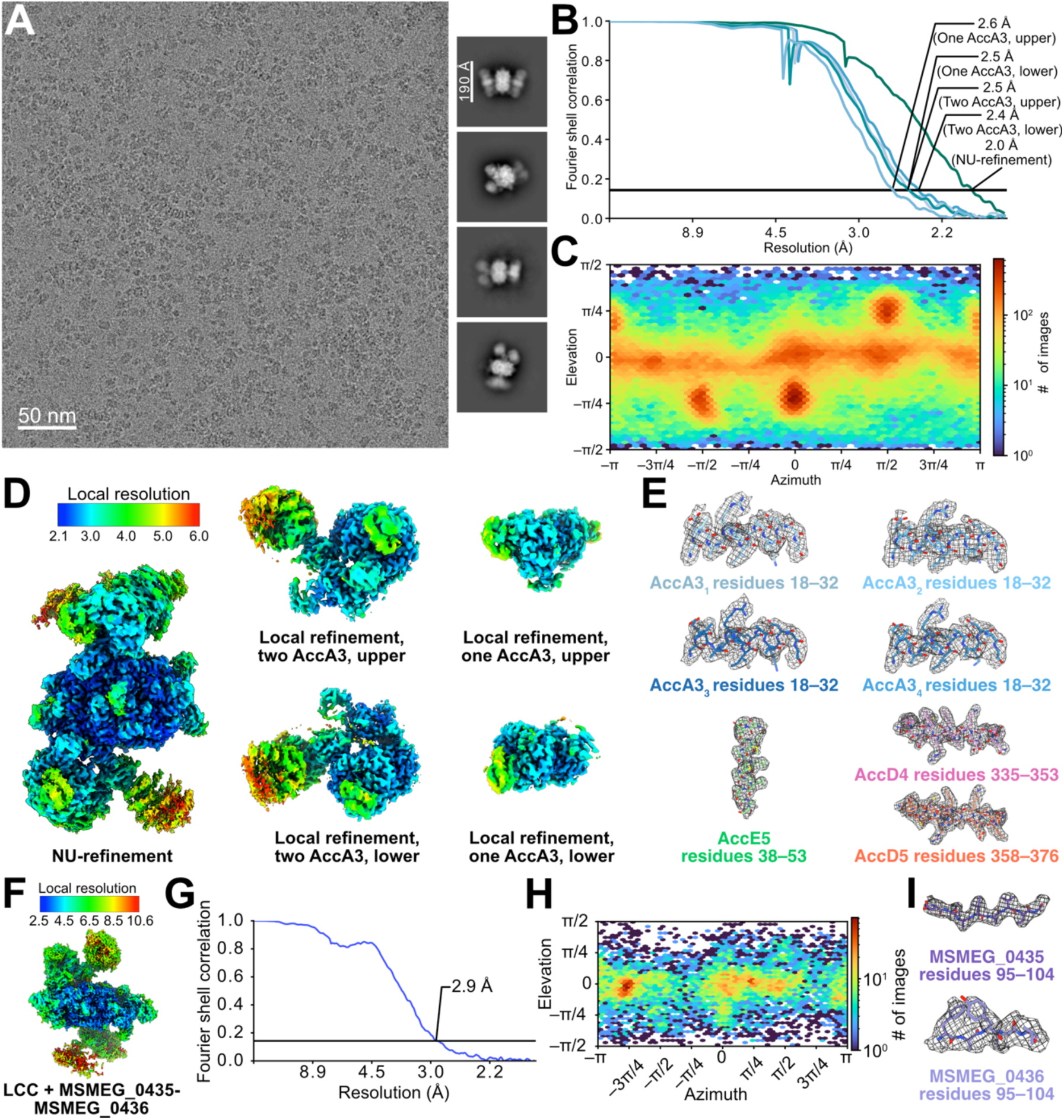
Cryo-EM map validation for the long-chain acyl-CoA carboxylase (LCC) complex with ATP, bicarbonate, arachidoyl-CoA, and propionyl-CoA. **A,** Example micrograph (*left*), and 2D class average images (*right*) showing the LCC complex in the presence of ATP, bicarbonate, arachidoyl-CoA, and propionyl-CoA. **B,** Fourier shell correlation curves following a gold-standard refinement and correction for the effects of masking for the overall non-uniform (NU) refinement, and with correction for the effects of masking for local refinement of BC domains from AccA3. **C,** Orientation distribution plot for particle images. **D,** Local resolution maps for the different regions of the LCC complex with ATP, bicarbonate, arachidoyl-CoA, and propionyl-CoA. **E,** Example model-in-map fit for selected regions of the LCC complex with ATP, bicarbonate, arachidoyl-CoA, and propionyl-CoA. **F,** Local resolution maps for the LCC complex with ATP, bicarbonate, arachidoyl-CoA, and propionyl-CoA, and with the MSMEG_0435-MSMEG_0436 complex bound. **G,** Fourier shell correlation curves following a gold-standard refinement and correction for the effects of masking for the LCC complex with the MSMEG_0425-MSMEG_0436 complex bound.

**Supplementary Figure 10.**
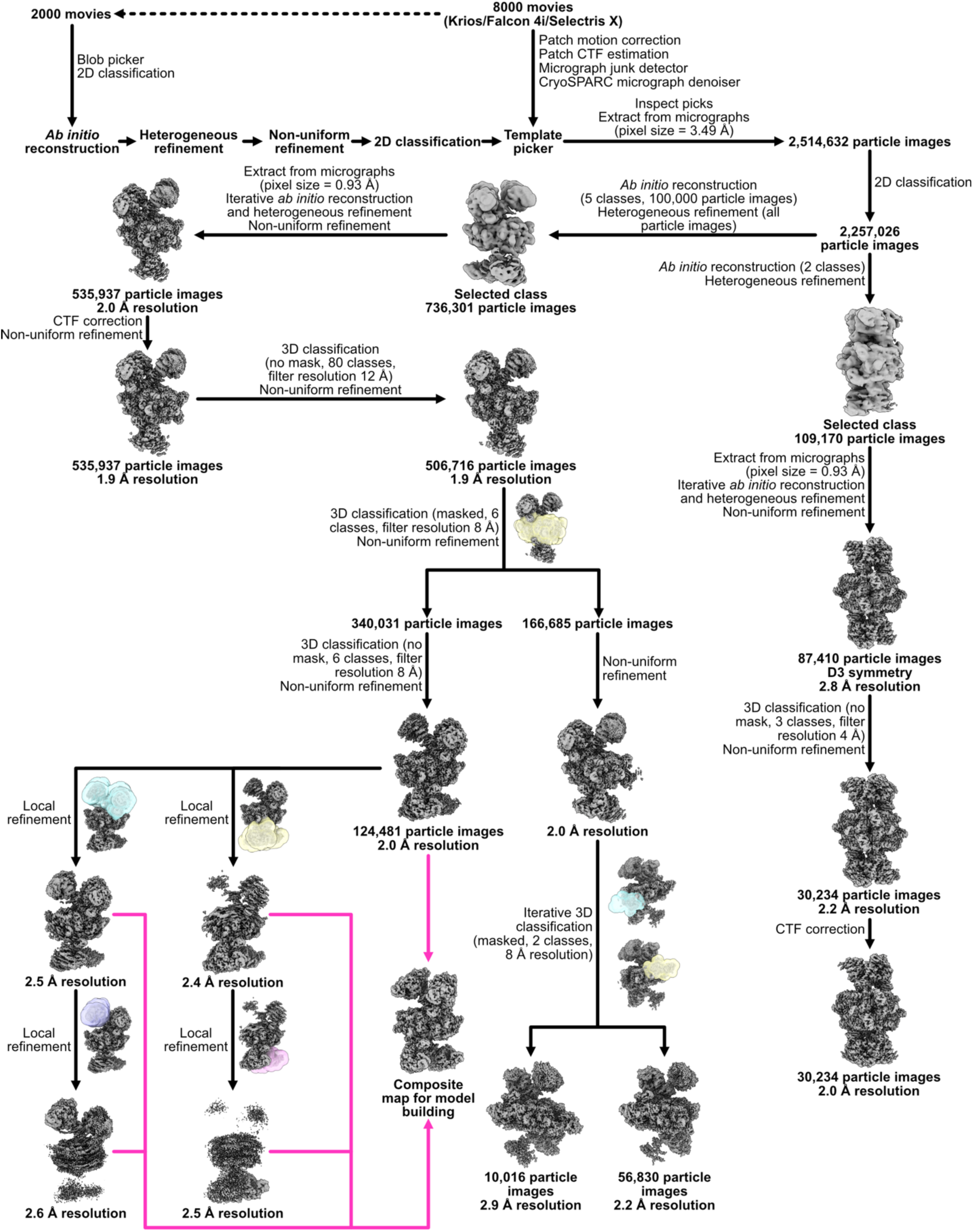
Workflow for calculating maps of the long-chain acyl-CoA carboxylase (LCC) complex with ATP, bicarbonate, arachidoyl-CoA, and propionyl-CoA.

